# Bringing the Ends together: Cryo-EM structures of mycobacterial Ku in complex with DNA define its role in NHEJ synapsis

**DOI:** 10.1101/2025.04.28.650930

**Authors:** Joydeep Baral, Ching-Seng Ang, Paul James McMillan, Kalyan Shobhana, Ayushi Saini, Elizabeth Hinde, Amit Kumar Das, Isabelle Rouiller

**Author notes:** Corresponding authors: (I.Rouiller) - *For Correspondence* (A.K.Das) (E.Hinde).

## Abstract

Non-homologous end joining (NHEJ) is the sole pathway for repairing double-strand breaks in *Mycobacterium tuberculosis* during dormancy, relying on mycobacterial Ku (mKu) and ligase D, with mKu as the rate-limiting factor. Despite its essential role, the lack of structural information on prokaryotic Ku has hindered understanding of the molecular mechanisms underlying bacterial two-component NHEJ machinery. Here, we present the first cryo-EM structures of mKu in DNA-bound and higher-order super-complex forms, revealing a Ku-mediated DNA synapsis mechanism unique to prokaryotes. Integrating cryo-EM with hydrogen-deuterium exchange mass spectrometry, we define key mKu-mKu dimerization, DNA-binding, and synapsis interactions essential for efficient NHEJ, bridging structure with function. Structure-guided *in silico* mutagenesis, coupled with electrophoretic mobility shift assays, identifies residues essential for DNA binding and synaptic assembly, which are crucial for NHEJ. Förster resonance energy transfer confirms DNA-dependent mKu oligomerization in solution, while live-cell imaging captures its spatiotemporal dynamics during DSB repair. These findings provide fundamental insights into the architecture and function of prokaryotic NHEJ, positioning mKu as a potential therapeutic target against tuberculosis and offering a framework for understanding DNA repair across bacterial species.

## Introduction

*Mycobacterium tuberculosis* (Mtb), the causative agent of tuberculosis, encounters constant threats to its genomic integrity from both environmental stressors and intrinsic factors (1). As an intracellular pathogen, Mtb resides within host macrophages in a dormant, non-replicating state (2). During dormancy, the pathogen is exposed to genotoxic agents such as reactive oxygen and nitrogen species, leading to substantial DNA damage, particularly double-stranded DNA breaks (DSBs) (3). DSBs are one of the most deleterious forms of genetic damage, as they threaten genome stability and cell viability. Consequently, efficient DNA repair is essential for Mtb’s survival in the hostile host environment (3).

Among the available repair mechanisms, the non-homologous end joining (NHEJ) pathway is uniquely capable of repairing DSBs in non-replicating cells (4). Unlike homologous recombination, which requires a template, NHEJ operates in a template-independent manner, making it indispensable during dormancy. The ability to exploit this pathway as a therapeutic target is particularly compelling when paired with DSB-inducing agents, offering new avenues to combat drug-resistant tuberculosis (5).

Bacterial NHEJ is significantly simpler than its eukaryotic counterpart, relying on two core components: the Ku protein and ATP-dependent DNA ligase D (LigD) (6,7). Ku binds to DNA ends, stabilizing them and protecting them from exonucleolytic degradation (8). It subsequently recruits LigD via its C-terminal tail, anchoring it to the damaged site (9). Together, these proteins form a minimal repair apparatus capable of ligating broken DNA ends (6).

A key step in NHEJ is DNA synapsis—the alignment and stabilization of broken DNA ends to enable ligation (10). Without intervention, exposed DNA ends are highly flexible and diffuse freely in solution. While the eukaryotic NHEJ apparatus has been extensively studied, with detailed 3D structures of synaptic complexes available (11), the mechanisms underlying bacterial DNA synapsis remain poorly understood. Current knowledge is limited to the two-component bacterial system, leaving significant gaps in understanding how synapsis is achieved and whether additional factors are involved (12).

Our recent studies on mycobacterial Ku (mKu) have shown its ability to form higher-order oligomeric complexes in the presence of DNA (8). However, whether this oligomerization is sufficient to mediate DNA synapsis remains unclear. In eukaryotes, the Ku70/80 heterodimer primarily acts as a DNA end-binding protein, recruiting additional NHEJ components for repair (13,14). Unlike its eukaryotic counterpart, mKu may play a more versatile role, potentially mediating synapsis independently or in collaboration with unknown partners. This presumed ability to align flexible DNA ends efficiently would distinguish mKu from other Ku proteins (8). Understanding the molecular mechanisms of mKu-mediated synapsis could offer unique insights into bacterial DNA repair and identify potential therapeutic targets.

Despite its biological importance, the structural and mechanistic basis of bacterial NHEJ remains poorly understood. Efforts to crystallize mKu have been hindered by the high flexibility of its C-terminal region, a trait observed in other bacterial Ku proteins such as *Bacillus subtilis* Ku (15). Additionally, mKu’s tendency to oligomerize in the presence of linear DNA introduces structural heterogeneity, complicating crystallographic and other structural studies (8). The small molecular size of mKu (31 kDa) poses further challenges for cryo-electron microscopy (cryo-EM), as resolving smaller proteins (<100 kDa) at high resolution remains difficult (16). Overcoming these challenges is essential for elucidating the molecular mechanisms underlying bacterial NHEJ and leveraging this knowledge for therapeutic development.

In this study, we employed cryo-EM to resolve the structure of mKu in complex with DNA, as well as to visualize the higher-order mKu-DNA oligomer (mKu-DNA supercomplex), which demonstrates novel mKu-mediated DNA synapsis. Using structural modeling and *in silico* alanine mutations, we identified critical residues for DNA binding and synapsis. We further explored the effects of DNA end modifications, such as hairpin structures, on the formation and stability of the DNA-protein complex and the supercomplex. Hydrogen-deuterium exchange mass spectrometry (HDX-MS) confirmed key interactions necessary for the formation of the mKu-DNA complex and synaptic assembly. Förster resonance energy transfer (FRET) was used to evaluate the stability and dynamics of the synaptic assembly in solution. Finally, live-cell imaging of *Mycobacterium smegmatis* fostering mKu (Mtb) allowed us to visualize and quantify the dynamic recruitment of mKu to DNA damage sites during the early stages of the NHEJ repair pathway.

## Methods

### Recombinant protein expression and purification

The recombinant plasmid (pET16b) containing the gene encoding mKu (Rv0937c) was a generous gift from Prof. Aidan Doherty, University of Sussex (9). The mutant constructs of mKu (mKu-Ala and mKuΔ12–15) were commercially synthesised and cloned into pET16b (Genscript) under a lactose-inducible promoter. The mKu-Ala mutant was designed by substituting all residues identified as critical for DNA binding with alanine, whereas the mKuΔ12–15 mutant was generated by deleting residues 12–15, which are critical for synapsis. The sequence validation of mKu and its mutant constructs is shown in Supplementary Figure 1. The plasmids were transformed into *E. coli* BL21 (DE3) C43 cells, and the proteins were overexpressed using isopropyl β-D-1-thiogalactopyranoside (IPTG). To obtain homogeneous protein samples, affinity (IMAC), ion-exchange (IEX), and size-exclusion (SEC) chromatography were performed sequentially. The overexpression and purification strategy of mKu WT has been previously described in detail (8). The purification protocol for mutant constructs (mKu-Ala and mKuΔ12–15) was identical to that of wild-type mKu. Protein purity was assessed by analytical size-exclusion chromatography and SDS-PAGE (Supplementary Figure 2). The molecular weight of mKu WT was determined by mass photometry using a Refeyn TwoMP mass photometer (Supplementary Figure 2.c). The buffer compositions used during purification are listed in the Supplementary Table 1.

### Cryo-EM sample preparation and data collection

The DNA substrates were commercially synthesized with the following sequences: (i) linear 40bp long double-stranded DNA (dsDNA): 5’-CCCCCCTGTCGCCGCCGACGTCTGTGATATGGCGTTGTTG-3’ and 5’-CCCCCCTGTCGCCGCCGACGTCTGTGATATGGCGTTG TTG-3’ (IDT). (ii) Short hairpin DNA (shDNA) with 21bp: 5’-GTTTTTAGTTTATTGGGCG CG-3’ and 34bp: 5’-CGCGCCCAGCTTTCCCAGCTAATAAACTAAAAAC-3’ (PDB:1JEY) (17). The DNA substrates were annealed by heating at 95° C followed by stepwise cooling to 4° C in a thermal cycler machine (ThermoFisher Scientific). The annealing buffer contained 10nM Tris pH 7.5, 50mM NaCl, and 1mM EDTA. The DNA substrates (dsDNA & shDNA) and mKu were incubated at an equimolar concentration of 5 µM for 30 mins at 4 °C. The dimeric molecular weight of mKu with affinity tag (66.8 kDa) was considered for molarity calculations as the protein is known to exist and function in the dimeric form. For grid preparation, 200 hole 1.2µm/1.3µm holey carbon grids (ProSciTech, cat# GCF200CU-1.2-25) were glow discharged (Quorum Technologies) at 15mA under vacuum for 3 mins. The longer than standard (30s) glow discharge time yielded better particle distribution. 4 µl of the sample were applied to the freshly glow discharged grids, which were blotted for 4 s at a blot force of 1 and plunge frozen in liquid ethane with a Vitrobot Mark IV (ThermoFisher Scientific) operated at 4 °C and 100% humidity. Following blotting and plunge freezing, the grids were clipped and loaded in a grid cassette for automated grid loading. Initial screening of grids was performed in Talos Arctica TEM (Thermo Fisher Scientific), followed by data acquisition in Titan Krios G4 (ThermoFisher Scientific) using the Falcon 4 detector. Movies were acquired at a magnification of x105,000 with a pixel size of 0.833 Å. For the mKu-dsDNA complex, a total of 4,682 movies were collected at a total dose of 40 e^-^/Å^2^ spread equally over 40 frames. Movies were recorded over a defocus range of -1 to -2.5 µm. For the mKu-shDNA complex, a total of 6,151 movies were collected, and the data collection parameters were unchanged.

### Cryo-EM processing and model building

Image processing and primary map generation were done using CryoSPARC v4.5.3(18). Motion correction and dose weighting were performed using the Patch Motion Correction job. Contrast Transfer Function (CTF) correction of the dataset was achieved with the Patch CTF job. Following the motion and CTF correction, the images were manually curated based on CTF fit resolution, ice thickness, and average intensity thresholds. Particle picking was initially done using the blob-picker with min and max diameters of 50 and 200 Å with an overlap of 0.5x diameter. Particle-picking parameters were adjusted, and different box sizes (256, 420 & 512 Å) were used to select and extract particles (mKu-DNA supercomplex) comprising 3 mKu dimers bound to 3 shDNA or 1.5 dsDNA molecules. As an alternate strategy, smaller box sizes (256 Å) were used to pick individual mKu_2_-DNA complexes to later combine into a composite map. However, the resulting 2D classes (not reported) had lower quality due to a reduced signal-to-noise ratio caused by the small size of the mKu dimer (∼70 Å in diameter). Thus, the particle picking was performed on the supercomplex rather than the unit mKu-DNA complex.

For the mKu-hairpin supercomplex, 1.15 million particles were primarily picked and extracted with a box size of 512 pixels, followed by 2D class averaging (100 classes) and manual curation to extract the unique orientations using a select 2D class job. Despite shDNA being shorter than dsDNA, mKu-shDNA oligomers had a higher curvature, requiring a larger extraction box to prevent loss of data (Supplementary Figures 3 & 4). The selected 2D classes were used as templates for template-based picking. Three rounds of 2D classification were done with manual curation between each cycle to obtain a final set of 229,557 particles. The resultant particles were used to generate *ab initio* maps using the Ab-initio reconstruction job in CryoSPARC. Three *ab initio* maps were generated with 43K, 110K, and 76K particles, respectively. Class II was selected for having the highest number of particles and better completeness in terms of 3D orientation. The *ab initio* maps were refined using the non-uniform refinement job, resulting in a global resolution (GSFSC 0.143) of 6.3, 6.8, and 4.1 Å (Supplementary Figure 5).

For the mKu-linear DNA supercomplex, the same workflow was used as above, with only the difference in the particle extraction box size (420 pixels). A total of 826,378K particles were used for *ab initio* reconstruction. *Ab initio* maps were classified into three classes with 101K, 418K, and 306K particles. The non-uniform refinement of the generated classes resulted in the best global resolution (GSFSC 0.143) of 4.3, 3.9, and 3.8 Å (Supplementary Figure 5). Finally, the density maps of the super-complexes were improved via density modification using the Phenix Resolve CryoEM job(19) (Phenix)(20). The native maps were used to isolate the densities corresponding to unit mKu-DNA complexes by running a model-based density modification job (Phenix). The resultant dimensions of the EM maps were 111 x 115 x 99 pixels (mKu-shDNA) and 190 x 100 x 109 pixels (mKu-dsDNA).

The dimeric model of mKu was generated using the AlphaFold2 web server implementation. (21). The model for linear DNA was obtained from our previous study(8), and hairpin DNA was extracted from PDB: 1JEY(17). The atomic models were initially fitted (rigid body fitting) into the density-modified maps of the unit complexes with the Local EM fitting module from the Phenix bundle in ChimeraX. Once the unit complexes (linear and hairpin) were fitted into the map, additional copies of the mKu dimer and DNAs corresponding to the supercomplexes were added to the map using the Local EM fitting module in ChimeraX(22). Following the rigid body docking of the models to maps, iterative refinement was performed using Real-space refinement in Phenix. Outliers and atomic clashes were resolved manually in Coot(23), and simultaneous real-space refinements were performed in Phenix. Model quality was assessed using a rotamer, geometry, and density fit analysis in Coot. After the final refinement job, the local and global resolutions of the maps were determined by ResMap(24), Local resolution map (Phenix), 3DFSC (Cryosprac), and GSFSC (Cryosprac) (Supplementary Figure 5).

### Electrophoretic mobility shift assay

DNA-binding activity of wild-type mKu and mutant variants (mKu-Ala and mKuΔ12–15) was assessed by gel-based EMSA using a 5′–FAM–labelled DNA substrate (Sigma). The DNA probe was 29-bp double-stranded DNA with 65.5% GC content (random sequence chosen to approximate *M. tuberculosis* genomic composition). To prepare the fluorescent dsDNA, the FAM-labelled oligonucleotide was annealed with an equimolar, unlabelled complementary strand by heating to 95 °C and gradually cooling to 4 °C in a thermal cycler (Applied Biosystems). Binding reactions contained 5 nM annealed FAM-dsDNA and mKu proteins at the indicated concentrations (5 nM, 500 nM, or 1 µM) in EMSA binding buffer (25mM Tris-Cl, 100mM NaCl, 30mM KCl, 0.1mM EDTA/Ethylenediaminetetraacetic acid, 0.05% TritonX-100, 500µg BSA, 2% v/v Glycerol, 2mM DTT /Dithiothreitol, pH 7.5). Reactions were incubated at 25 °C for 30 min. Samples lacking protein served as negative (free-DNA) controls; reactions lacking DNA served as buffer controls. Complexes were resolved on 10% native polyacrylamide gels in 0.5× TBE (pH 7.5) at a constant 90 V for 3 h. Gels were imaged on a ChemiDoc MP system (Bio-Rad) using a fluorescein filter. Results were evaluated qualitatively based on the relative presence or absence of shifted bands, with free DNA migrating at the bottom of the gel and DNA–protein complexes appearing as slower-migrating bands higher in the gel.

### Hydrogen-deuterium exchange mass spectrometry

Hydrogen-deuterium exchange (HDX) labeling of the protein was conducted at 20°C. 3 µL aliquots of the purified protein (∼15 µM, dimer) were added to 55 µl of either non-deuterated (50 mM potassium phosphate buffer, pH 7.4 in H_2_O) or deuterated (50 mM potassium phosphate buffer, pH 7 in D_2_O) buffer and incubated for 0, 0.1, 1, 10 and 100 mins using a PAL Dual Head HDX Automation Manager (Trajan/LEAP) controlled by ChronosHDX software. At the end of the incubation time, the reactions were quenched by mixing 50 µL of the deuterated protein sample with 50 µL of quenching buffer (50 mM potassium phosphate buffer, pH 2.3, 4 M guanidine hydrochloride, and 0.1% n-Dodecylphosphocholine) at 1°C.

For online pepsin digestion, 80 µL of the quenched samples were then passed through an immobilized Enzymate BEH pepsin column (2.1 × 30 mm; Waters), equilibrated in 0.1% formic acid at 100 µL/min. To reduce peptide carryover, 1% w/w n-Octyl-β-d-glucopyranoside was added to the column wash solution (1.5 M guanidine hydrochloride, 4% acetonitrile, 0.8% formic acid). The proteolyzed peptides were captured and desalted using a VanGuard BEH C18 trap column (1.7 µm, 2.1 × 5 mm; Waters), then eluted with a gradient of acetonitrile and 0.1% formic acid (5% to 40% over 8 minutes, 40% to 95% over 0.5 minutes, held at 95% for 1.5 minutes) at a flow rate of 80 µL/min, and separated on an ACQUITY UPLC BEH C18 analytical column (1.7 µm, 1 × 50 mm; Waters), delivered by an ACQUITY UPLC I-Class Binary Solvent Manager (Waters).

Mass spectrometry was performed using a SYNAPT G2-Si mass spectrometer (Waters) equipped with ion mobility. Instrument settings were as follows: 3.0 kV capillary voltage, 40 V sampling cone, source temperature at 100°C, and desolvation temperature at 40°C. The desolvation gas flow was set to 800 L/hr, and the cone gas flow was 100 L/hr. The high-energy trap collision energy was ramped from 20 to 40 V. Mass spectra were acquired with a 0.4-second scan time and continuous lock mass (Leu-Enk, 556.2771 m/z) for mass accuracy correction. Data were collected in HDMSE (ion mobility) mode, and peptides from non-deuterated samples were identified using Protein Lynx Global Server (PLGS) v3.0 (Waters). Peptide selection stringency was ensured by applying additional filters: 0.3 fragments per residue, minimum intensity of 5000, maximum MH+ error of 5 ppm, retention time RSD of 10%, and file threshold of 4 out of 6 HDMSE files. Deuterium uptake values were calculated for each peptide using DynamX 3.0 (Waters). Deuterium exchange experiments were conducted in triplicate for each time point. Peptides with statistically significant differences in HDX were identified using Deuteros 2.0 software (25), employing a hybrid significance test with a 99% confidence interval.

### Förster Resonance Energy Transfer

Single-stranded DNA molecules were commercially synthesized (IDT) and annealed to get the blunt-ended dsDNA substrate. Individual DNA stands contained- Cy3 as a donor and Cy5 as an acceptor. The following sequences correspond to the two complementary strands of the DNA substrate: 5’Cy5/CCACACTGTCGCCGCCGACGTCTGTGATATGGCGTTGTTGCGCCG CCGAC/3’Cy3Sp and 5’Cy3/GTCGGCGGCGCAACAACGCCATATCACAGACGTCGGCGGCGACAGTGTGG/3’Cy5Sp to have a donor and acceptor fluorophores at opposite ends. The fluorescence signal was detected at λ_excitation_ of 554 nm (λ_ex_ of Cy3) and λ_emission_ of 670 nm (λ_em_ of Cy5) using Cary Eclipse Fluorescence Spectrometer (Agilent). The negative control set with no mKu was prepared with 250nM DNA in 10 mM PBS buffer pH 7.4. Buffer control was used as blank to remove the background signal from the buffer. For the FRET assay, a fixed concentration of 250 nM DNA solution was titrated with increasing concentrations of mKu (0-305 nM) until the fluorescence emission signal reached saturation. Incubation time between injection and data acquisition was kept constant at 1 min. The experiment was performed in triplicates at 25 °C using a Peltier system attached to the Spectro fluorimeter. Wave scans (emission: 570-700nm) were performed for respective mKu concentrations to quantify the fluorescence intensities at λ_emission_ maximum (Supplementary Figure 6). Finally, the normalized fluorescence intensities (λ_max_) corresponding to respective concentration mKu were graphically represented (Fig 5.d).

### Live cell confocal microscopy

The *Mycobacterium smegmatis* mc² 155 *Δku ΔligD* strain (Mgm153) (26) was kindly provided by Prof. Michael S. Glickman (Memorial Sloan Kettering Cancer Center, New York). This strain is deficient in non-homologous end joining (NHEJ) factors, with the intrinsic *ku* and *ligD* genes deleted. Cells were cultured in Middlebrook 7H9 medium supplemented with 0.5% dextrose, 0.5% glycerol, 0.05% Tween 80, and OADC (oleic acid, bovine albumin, sodium chloride, dextrose, and catalase). To prevent contamination, 100 µg/ml ampicillin was added. Where required, 50 µg/ml hygromycin B was included for selection. The plasmid pYUB1062-GFP (27) (Plasmid #117695, Addgene) was used to generate the mKu–GFP fusion construct (Genscript). The recombinant plasmid was electroporated into electrocompetent Mgm153 cells using a MicroPulser electroporator (Bio-Rad) with one pulse at 2.5 kV, EC2 setting, in a 0.2 cm cuvette. Transformed cells were recovered in Middlebrook 7H9 medium and plated onto Middlebrook 7H10 agar containing hygromycin B, followed by incubation for 72 h. For imaging, mKu–GFP–expressing Mgm153 and control cells were cultured in liquid medium to an OD₂₆₀ of 0.6. Expression was induced with 0.2% acetamide at 37 °C for 20 h. Cells (1 ml) were harvested and stained with 0.5% (v/v) Hoechst 33342 (ThermoFisher) for 30 min. The stained suspension was placed on a 1% agarose pad mounted on a glass slide and maintained at 37 °C in a temperature- and humidity-controlled chamber. Fluorescence imaging was performed using a Zeiss LSM900 inverted confocal microscope equipped with an Airyscan 2 detector and a 63× oil immersion objective (working distance: 140 µm). Images were acquired in laser scanning confocal mode, with each dataset consisting of 58 image stacks captured at 5.65 s intervals over 300 s. A total of five biological replicates were acquired for this study. Laser-induced DNA damage was introduced using a 405 nm laser targeted to a 1.0 × 0.8 µm region of interest (ROI) at 5 mW laser power and 100% bleach intensity. Damage was applied after the third frame (11.3 s), with frames 1–3 (0–11.3 s) serving as pre-damage references, and frames 4–70 (16.96–300 s) capturing post-damage responses. For control experiments, no laser damage was applied, while acquisition parameters and ROI dimensions remained identical. DNA was visualized using the DAPI channel (400–500 nm detection range), and GFP-tagged mKu was imaged in the GFP channel (495–600 nm). Image stacks were processed in FIJI (28). Fluorescence intensity fluctuations of Hoechst 33342 and mKu-GFP at the ROI were quantified across the 58 frames using the Plot Z-axis profile function (FIJI). The recorded intensities were normalized using the following equation: 𝐼(𝑁𝑜𝑟𝑚) = 𝐼𝑛 − 𝐼𝑚𝑖𝑛/(𝐼𝑚𝑎𝑥 − 𝐼𝑚𝑖𝑛). Upon normalization, the normalized intensity was plotted as a function of time along with the standard error of the mean (SE). For each time point, the normalized fluorescence intensities of Hoechst 33342 and mKu-GFP at the laser-damaged ROI were divided by those at the undamaged ROI. These damaged-to-undamaged ratios were plotted over time and analyzed for statistical significance. Data statistics and normalizations were performed in Origin(Pro), Version 2024. The raw data and biological replicates are provided for reference (Supplementary Figure 7 and Supplementary file).

## Results

### Mycobacterial Ku homodimers form higher order oligomers in complex with linear and short hairpin DNA molecules

Our previous study demonstrated that mycobacterial Ku (mKu) forms higher-order oligomers in the presence of DNA (8). These oligomeric complexes were obtained using chemically synthesized 40 base pairs long linear dsDNA molecule. However, the resultant complexes were of heterogeneous lengths, measuring more than 200 nm, corresponding to several dozens of mKu dimers (∼6 nm each) adjacently placed, forming a filamentous structure (Supplementary Figure 3).

In an attempt to obtain structurally homogenous complexes of DNA and mKu, we assembled DNA-mKu complexes using a hairpin-DNA molecule inspired by the human Ku70/80-DNA complex crystal structure (PDB: 1JEY). This structure used a short-hairpin DNA (shDNA) comprising a 34-mer and a 21-mer ssDNA, annealed to form dsDNA with a hairpin at one terminus. We predicted that the bulky terminal hairpin would prevent mKu from forming higher-order oligomers. However, contrary to our expectations, we observed long filamentous oligomers (Supplementary Figure 3) approximately 6 nm in diameter and ∼50-100 nm in length rather than well-defined or discrete particles formed by a mKu dimer and DNA. The mKu-shDNA particles were shorter with more curvature than those obtained with dsDNA, indicating that the bulky hairpin inflicted some hindrance in the system but was insufficient to prevent assembly into filaments (Supplementary Figure 3). Importantly, the inherent propensity of mKu to form filaments in the presence of DNA did not impede cryo-EM structure determination, as the observed heterogeneity was limited to filament length, reflecting variations in the number of mKu dimers per filament. This enabled robust structural reconstruction of the mKu–DNA unit complex as well as its higher-order oligomeric assembly (mKu–DNA supercomplex).

### Cryo-EM structures of mKu homodimer in complex with dsDNA and shDNA molecules

To determine the structure of the mKu-DNA supercomplex, 5 µM of the purified mKu dimer was incubated with an equimolar concentration of dsDNA or shDNA. The 3D reconstruction was performed using the single-particle cryo-EM image analysis with box sizes encompassing 3 mKu dimers and 1.5 dsDNA molecules or 3 shDNA. These box sizes were chosen based on a series of 2D classifications. For clarity, the following structural nomenclature is used throughout: a unit complex refers to an mKu homodimer bound to a single dsDNA or shDNA molecule at a 1:1 stoichiometry; a supercomplex designates higher-order oligomers formed through the assembly of multiple unit DNA–protein complexes (Figure 1). The synaptic complex represents the simplest form of supercomplex, consisting of two mKu dimers and two dsDNA molecules.

**Figure 1.**
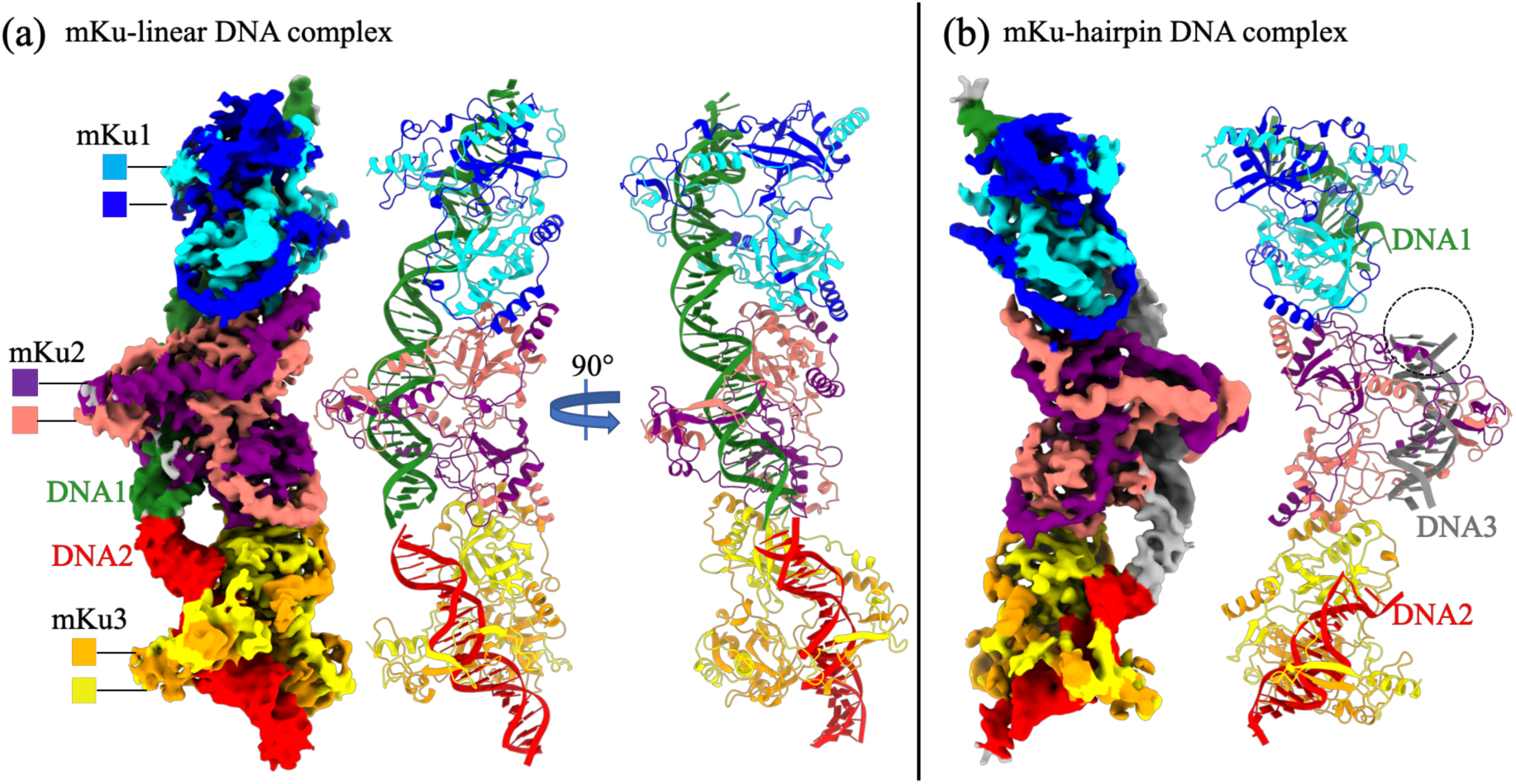
**| Cryo-EM structures of the prokaryotic Ku-DNA complex** (a) EM map and structural model of the mKu-linear DNA supercomplex. Three adjacent mKu dimers are bound to two linear DNA molecules, with each protein and DNA chain colour-coded as shown. The C-terminal 46 residues of the mKu protein are not visible due to missing electron density. (b) EM map and structural model of the mKu-hairpin DNA supercomplex. Three adjacent mKu dimers are bound to three DNA molecules, having a terminal hairpin. The hairpin region (dotted circle) is unresolved due to missing electron density.

Our maps had a GSFSC (gold-standard Fourier shell correlation) resolution of 3.8 Å for dsDNA and 4.1 Å for shDNA. The resolutions of these maps were sufficient to enable us to build several models (Figure 1). The data collection, validation, and refinement statistics are provided in Supplementary Table 2. The cryoEM workflow, map validation, and resolution estimation are presented in Supplementary Figures 4,5. These maps showed clearly defined densities for the Ku dimers and the DNA spanning the dimers. Each Ku dimer has an approximately triangular prism 3D shape with a hole at the top of its lateral face through which DNA molecules are threaded. The two Ku molecules are highly intertwined within the dimer. The Ku dimers interact extensively at the bottom of their lateral faces at a defined angle, forming a continuous protein filament. However, the flexible C-terminal tail of mKu could not be traced on the EM map because of the lack of corresponding density. The Continuous densities for the DNA molecules are resolved in the EM maps, with the major and minor grooves of the DNA clearly visible. The DNA molecules are better resolved in the linear DNA density map and are positioned end-to-end along the central axis of the Ku filament. Continuous density for DNA is also present in the EM map. The density in the shDNA was less resolved with no density was observed for the hairpin itself, suggests that adjacent shDNA molecules were in close proximity but in an energetically unfavourable arrangement, possibly due to hindrance from the bulky hairpin moiety.

### Molecular hand-clenching facilitates mKu dimerization

Mycobacterial Ku is a dimer in solution and partially resistant to SDS(8). The melting temperature of the dimer is 50° C, which is considerably high for a protein in its apo state(8). Our models of mKu dimer (PDB: 8V53,8VF5) reveal that the interaction between the two monomers of mKu resembles two human hands clenched together (Figure 2.a, b). The regions of contact in the dimeric interface are broadly divided as (i) the bridge (residue: 23-91), (ii) the pillar (α-helix-#4), and (iii) the sling (residue: 167-227) comprising a helix-turn-helix (HTH) motif (Figure 2.c). The bridge comprises three beta-hairpins (ß2 – ß4) in antiparallel contact with the opposite monomer, mimicking crisscrossed human fingers (numbered 1-3, shown in Figure 2). Building on the observations from EM structures, HDX-MS revealed that β-hairpins 2 and 3 were protected from deuterium-to-hydrogen exchange (Figure 3. a, b). Such protection in HDX-MS indicates that these regions are involved in interactions, with the residues being either fully or partially buried within the protein dimer. Our models suggest total of 20 H-bonded contacts and two salt bridges (Ku1-E65:R30-Ku2 & Ku2-E65:H32-Ku1) interactions in the bridge region, as calculated from the dimer structure using LigPlot+ (Supplementary Figure 8 and Table 3). The pillar and β-hairpin4 of Ku1 form a cleft that houses Ku2 β-hairpin2 via non-bonded contacts (Figure 2. c. ii). Finally, the sling arises from two respective monomers and wraps around the whole complex in opposite directions, enhancing the compactness of the dimer. The sling forms significant contact with the β-barrel of the opposite monomer (Figure 2.c.iii). Overall, these three interfaces (bridge, pillar & sling) contribute to the relative rigidity and stability of the mKu dimer.

**Figure 2.**
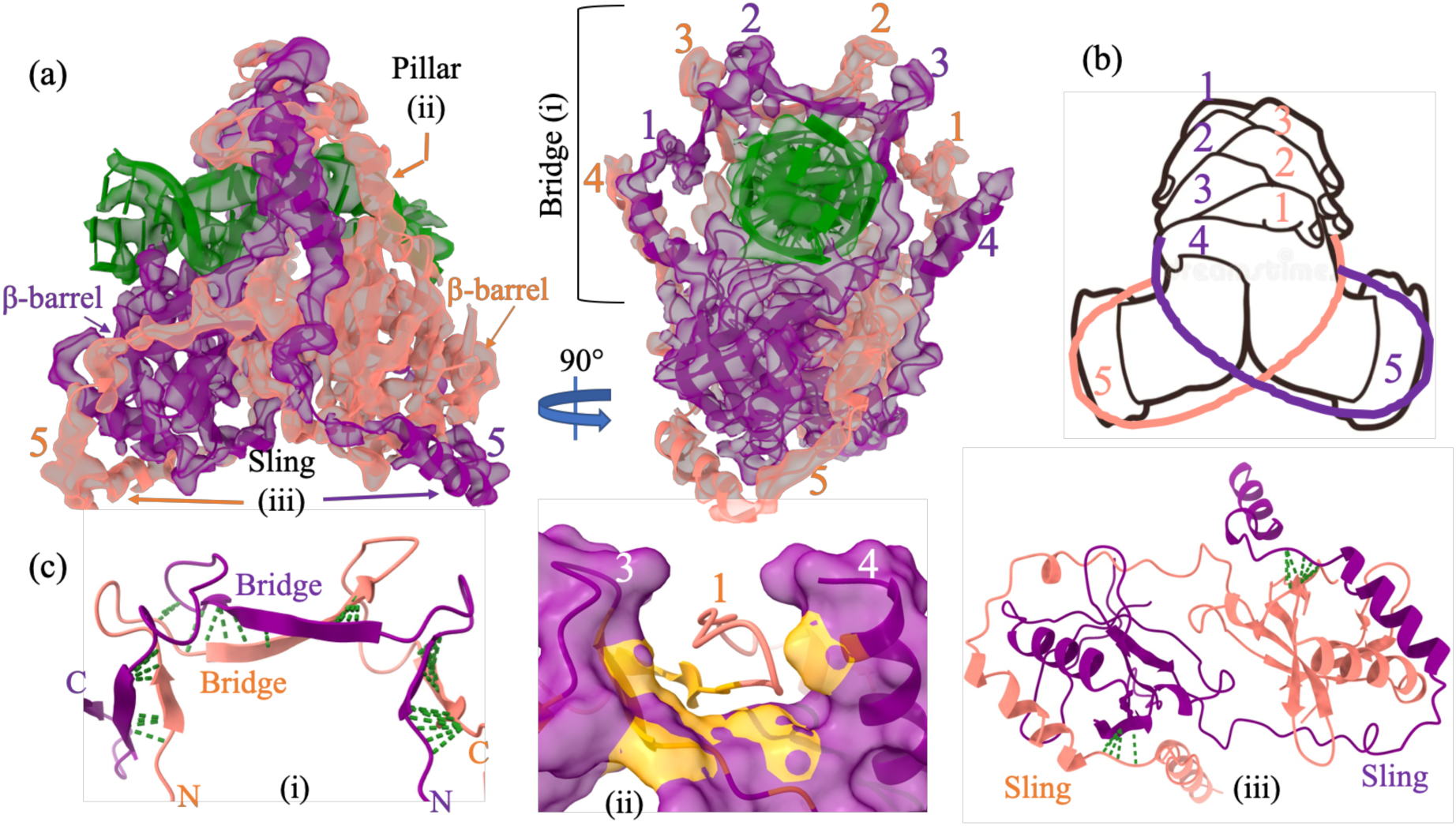
**| Molecular hand-clenching of mKu homodimer** (a) EM map with the overlaid model showing two mKu monomers (coloured purple and salmon) interacting through three regions: (i) Bridge (1–3), (ii) Pillar (4), and (iii) Sling HTH motif (5). The numeric notations (1–5) denote the contact regions and do not indicate the protein’s secondary structural features, such as β-hairpins or helices. (b) A schematic representation highlights the resemblance between the mKu dimer and clenched human hands, where the fingers correspond to the contact regions (1–5). The outline image of human hands was obtained from Deamstime.com (c) Detailed views of the mKu dimer interfaces (i, ii, and iii). For clarity, the dimeric complex is truncated to enhance visualization. Green dotted lines represent the contact points.

**Figure 3.**
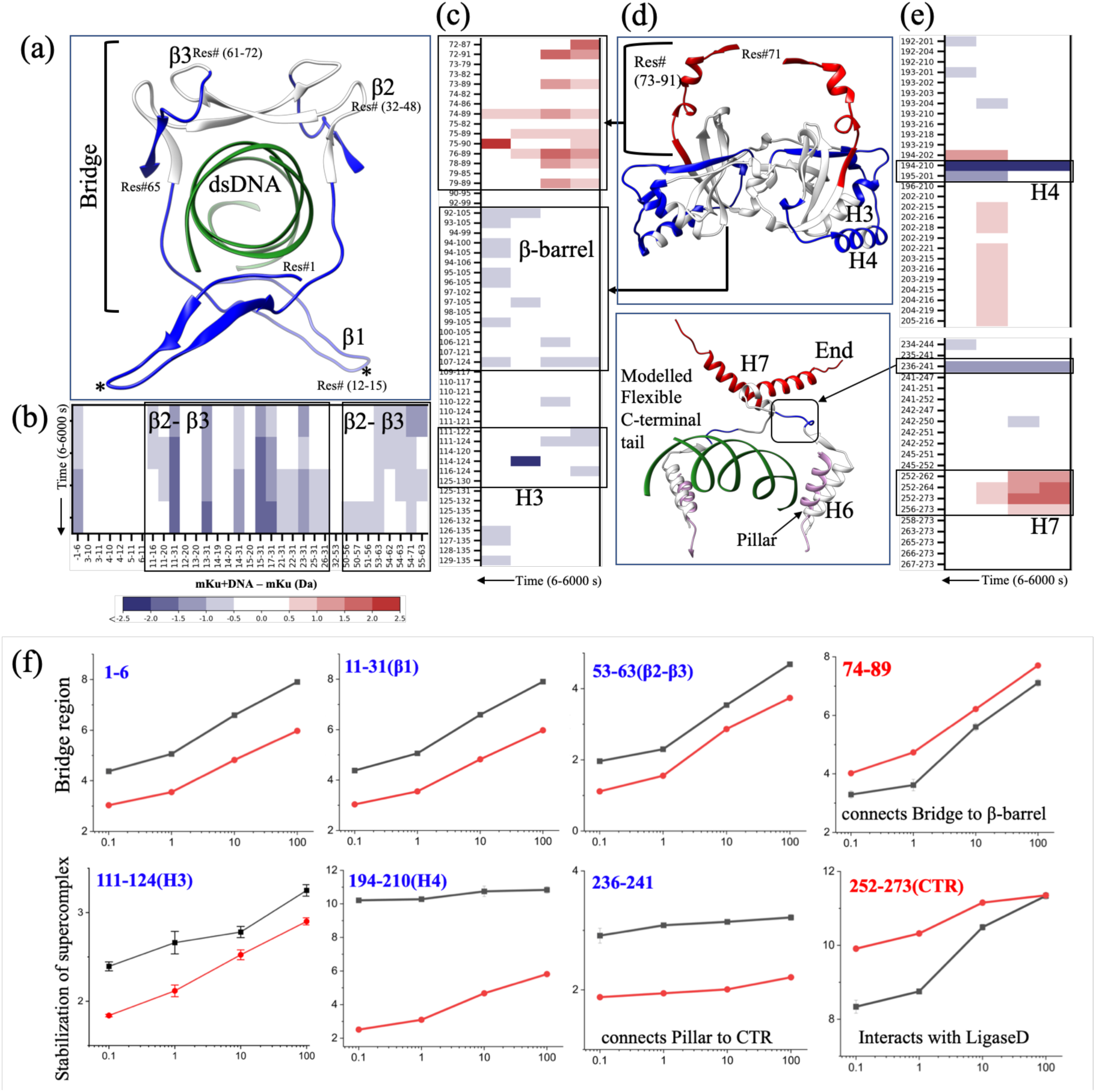
**| HDX-MS analysis of protein-protein and DNA-protein interactions in the mKu and mKu-dsDNA complex** (a & d) Truncated model structures are colour-coded based on relative deuterium uptake: protected regions are shown in blue, while deprotected regions are depicted in red. The relative uptake represents the difference in D₂O incorporation (mKu-DNA minus mKu) for respective peptide stretches. (b, c & e) Chicklet plots illustrate global, time-dependent uptake profiles across 0.1–100 minutes. (f) Comparative local uptake plots highlight critical regions, with mKu-DNA in black and apo mKu in red. Plot labels are coloured blue or red, indicating protected or deprotected nature.

### Identification and cross-validation of critical residues in mKu-DNA complex

Ku homodimer interacts with the DNA via the central pore located at the center of the dimer. The DNA binding pore is lined with positively charged residues (Supplementary Figure 9), perfectly suitable for a sequence non-specific interaction with the sugar-phosphate backbone of a dsDNA molecule. In the mKu-DNA complex, the DNA acts like a nut, sliding into the DNA binding pore of mKu, similarly resembling a bolt. This mode of nut-bolt assembly of Ku and DNA is conserved across species(7). However, the functional aspects of Ku vary in lower and higher organisms due to the presence or absence of additional domains at the N and C termini of the protein(7).

The DNA-mKu interaction is mediated majorly by positively charged residues (Arg & Lys) of the mKu dimer and the negatively charged sugar-phosphate backbone of the DNA (Figure 4). mKu interacts with the DNA at two distinct regions on the DNA, at the DNA termini (End) and the central area of the DNA (Mid) encompassed by the bridge region of mKu (Figure 4). The residue-specific contacts are mapped in a 2D figure for better visualization (Figure 4.b). Residues Y(21 & 44), Q33, and R(43,46,58,62 & 88) were found to form H-bonded contacts with the backbone of the DNA double-helix (Figure 4.b). Similarly, the stretch of interacting residues lining the DNA binding pore was found protected in HDX-MS (Figure 3.a, b). In addition to the H-bonded contacts, several non-bonded contacts stabilize the DNA-Ku complex. Overall, the tight binding of the DNA at the positively charged pore of mKu explains the previously reported high thermal stability of the mKu-DNA complex (Tm = 70° C)(8).

**Figure 4.**
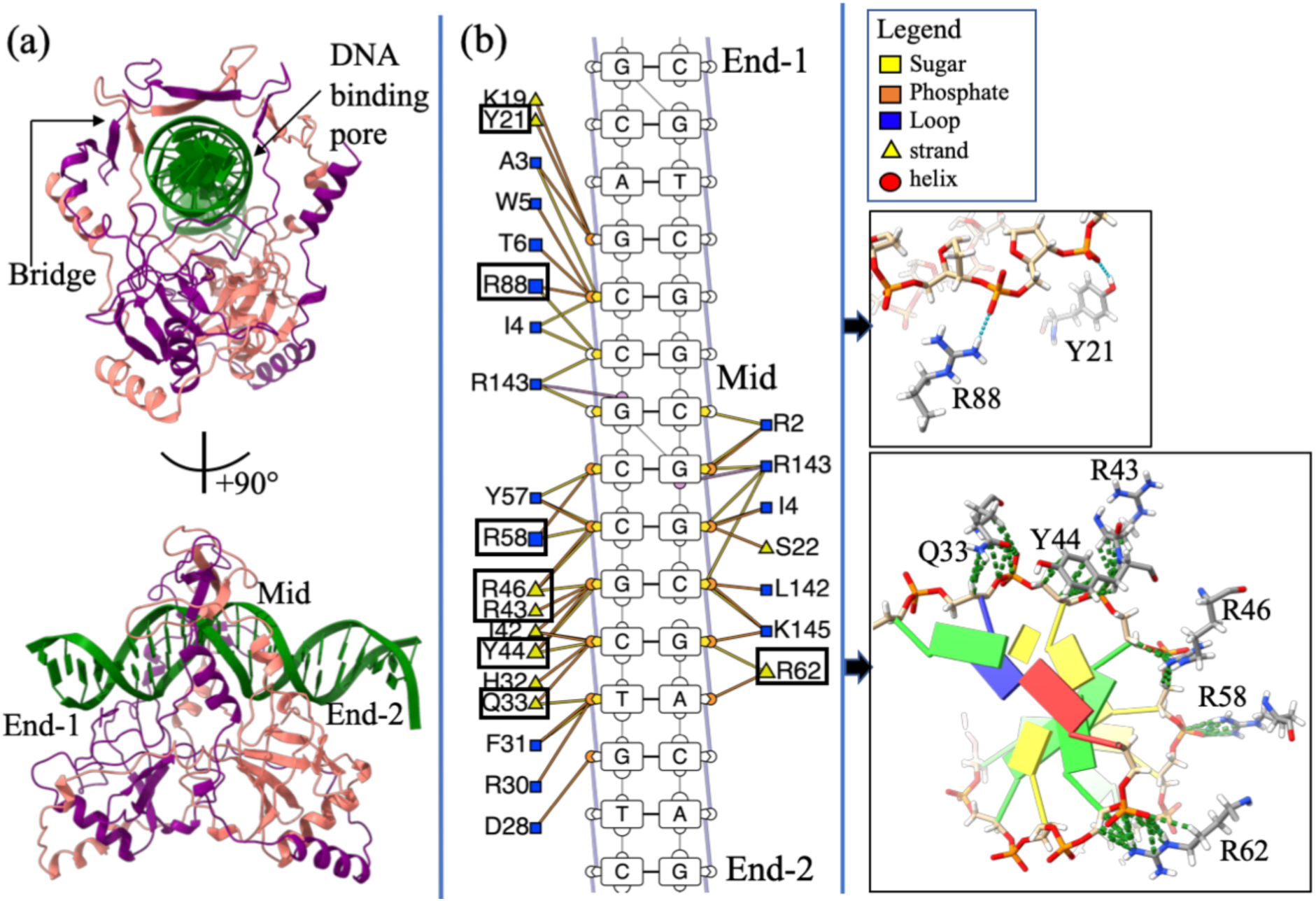
| **DNA-mKu interactions** (a) Structural representation of the unit dimeric mKu-DNA complex, highlighting the DNA-binding pore of mKu and the regions of DNA contact. (b) 2D contact map illustrating DNA-protein interactions, with critical contacts detailed in the adjacent panel. Key contacts are indicated by green dotted lines.

To identify residues essential for DNA binding, we performed *in silico* alanine scanning of the mKu–DNA interface and compared the stability of mutant complexes to the wild type. Mutations at Y21 and R(58,88) moderately destabilized the DNA–protein complex, whereas substitutions at Y44, Q33, and R(43,46,62) were highly destabilizing (Supplementary Table 4). AlphaFold3 models of the mKu-Ala–DNA complex and synaptic super-complex corroborated these predictions, showing neutralization of the positively charged lining of the DNA-binding pore and partial accommodation of DNA within the channel (Figure 6.a,b). The mKu-Ala variant was generated by substituting all the identified DNA-binding residues (Supplementary Table 4) with alanine. Notably, the AlphaFold3 model of mKu-Ala did not predict disruption of DNA synapsis *in silico*. To experimentally validate these predictions, we compared the DNA-binding capacity of wild-type (WT) mKu and the alanine-substituted mutant (mKu-Ala) using electrophoretic mobility shift assays (EMSA). Alanine substitutions caused a marked reduction in DNA-binding affinity (Figure 6.e,f). At equimolar concentrations with FAM-labeled DNA (5 nM), mKu-WT completely shifted the free DNA, whereas the mKu-Ala mutant showed no detectable binding, as indicated by the persistence of the free DNA band and absence of higher-order DNA–protein complexes (Figure 6.e). Even at a 200-fold molar excess of mKu-Ala (1 µM), only limited depletion of free DNA and weak complex formation were observed (Figure 6.f).

**Figure 5.**
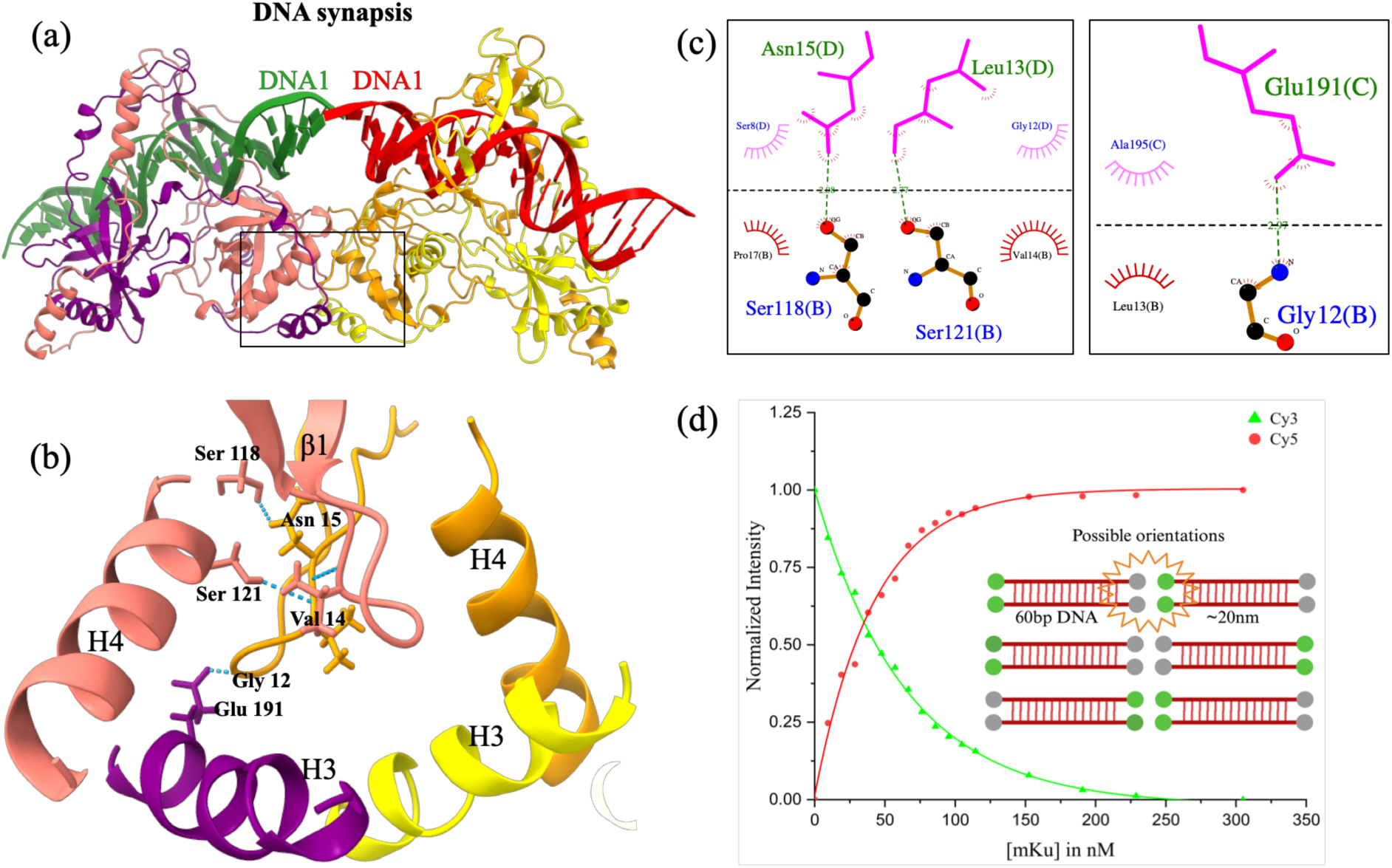
**| Higher order oligomerization of mKu facilitates DNA synapsis** (a) Cryo-EM structure of the mycobacterial Ku–DNA synaptic complex, with individual protein chains shown in distinct colours. (b) Close-up view of the β-hairpin1 interface, highlighting residues 12–15 (GLVN) critical for oligomerization. (c) Schematic 2D representation of atomic contacts between adjacent mKu homodimers that stabilize the mKu–DNA supercomplex. Chains a–d correspond to individual mKu monomers. (d) FRET assay demonstrating that wild-type mKu promotes DNA synapsis by juxtaposing broken DNA ends. Normalized donor (Cy3, green) and acceptor (Cy5, red) fluorescence intensities are shown across increasing mKu concentrations (0–305 nM).

**Figure 6.**
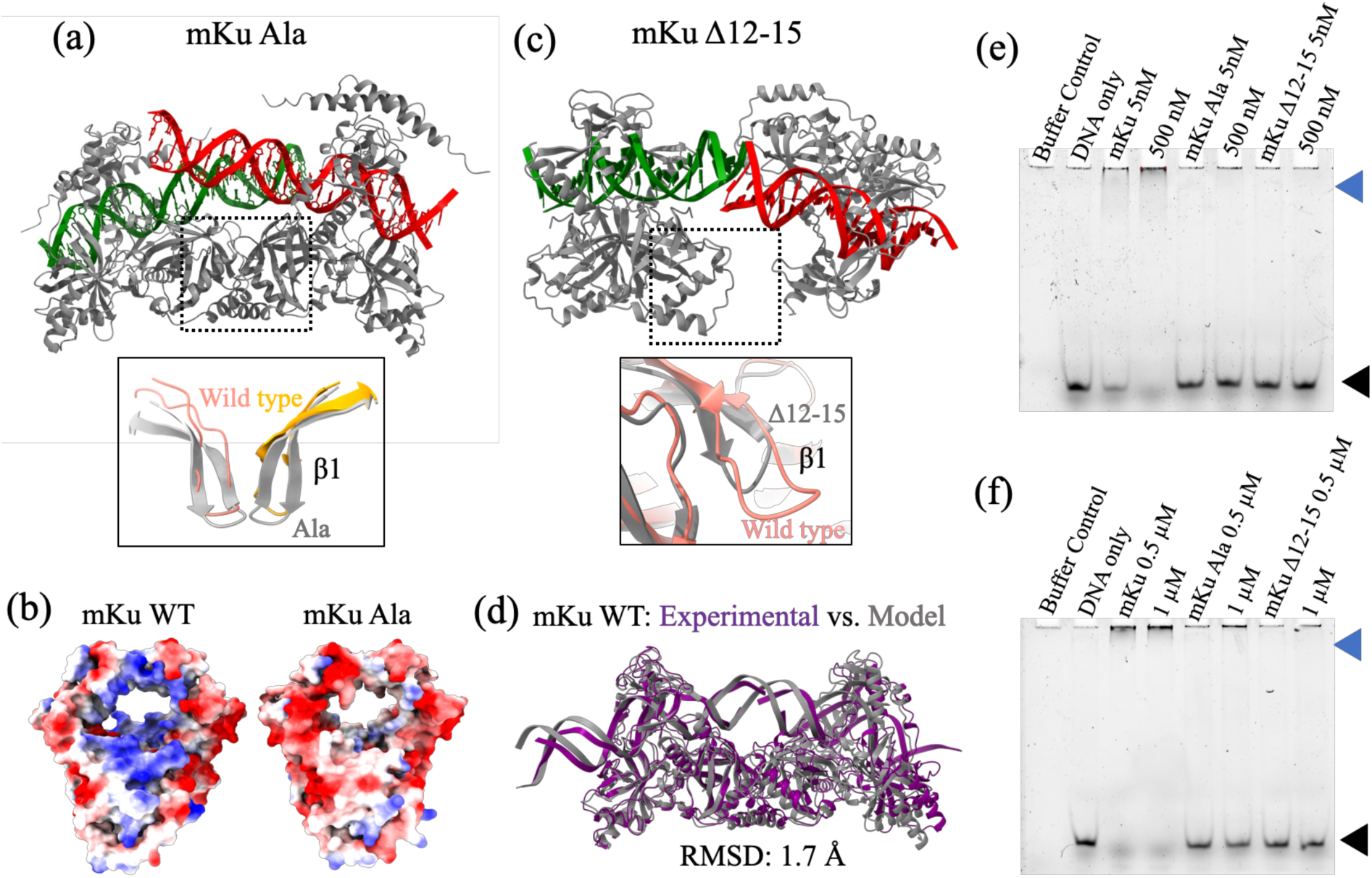
**| Mutations in the central pore and β-hairpin1 impair DNA binding and synapsis** (a) Modelled supercomplex of the mKu Ala mutant, with a zoomed-in view of the synaptic interface showing the β-hairpin1 region compared to wild-type (WT) mKu. (b) Coulombic electrostatic surface potentials of mKu WT (cryo-EM structure) and mKu Ala (AlphaFold3 model). The central pore’s positive charge in WT is neutralized in the mutant dimer. Blue to red colour scale represents positive to negative charge. (c) Modelled supercomplex of the mKu Δ12–15 mutant, showing disruption of the synaptic interface due to the shortened β-hairpin1, highlighted in the zoomed-in view. (d) Comparison of the experimental cryo-EM structure of the mKu–DNA synaptic complex with the corresponding AlphaFold3 model. (e, f) Comparative EMSA assays showing DNA binding by mKu WT and mutants (mKu Ala and mKu Δ12–15). Free DNA is indicated by a black arrow, and DNA–protein complexes by a blue arrow.

### Higher order oligomerization of mKu directs DNA synapsis

DNA end binding of Ku has been proposed to be the rate-limiting step in NHEJ(6). Our data suggests that mycobacterial Ku has a critical role in initiating the DSB repair. In addition to DNA binding and protecting broken DNA ends from exonucleases(8), mycobacterial Ku independently facilitates DNA synapsis via mKu dimer oligomerization (PDB: 8VF2, 8VF4). The mKu-mediated synapsis is formed by the simultaneous association of the mKu dimer-1 (chain A & B) bound to DNA1 (chains E & F) with an adjacent mKu dimer-2 (chain C & D) bound to DNA2 (chains G & H). The DNA-mKu interactions are discussed in detail in the preceding section. In the mKu1-mKu2 super-complex interface, we found that ß-hairpin1 from the adjacent mKu dimers make hydrogen-bonded and non-bonded contacts. Specifically, residues 12 to 15 (GLVN) present in the ß-turn region of the ß-hairpin1 motif were found to interact with the ß-hairpin1 motif of the adjacent mKu dimer (mKu2) and the surrounding helix-3 and 4 (Figure 5). The helix 3 and 4 from the adjacent mKu dimers (mKu1 & 2) create a four-helix pocket that creates and stabilizes the aforementioned super-complex interface. The following H-bonded contacts were observed between ß-hairpin1 and helix 3,4: mKu1-B-12-Gly to mKu2-C-191-Glu (Bond distance: 2.97 Å), mKu1-B-118-Ser to mKu2-D-15-Asn (Bond distance: 2.08 Å), mKu1-B-121-Ser to mKu2-D-13-Leu (Bond distance: 2.77 Å). Here, the residues are annotated as Dimer ID-Chain ID-Residue Number-Residue Name. To assess the contribution of the residues mentioned above in ß-hairpin1, we have modeled the mutants of mKu using AlphaFold3. We found that the deletion mutants- mKuΔ12-15 and mKuΔ1-15-resulted in complete disorientation and disassembly of the synaptic complex (Figure 5.d & Supplementary Table 5). In contrast, alanine substitution of 12-15 did not impact the complex assembly. Functional assessment by EMSA confirmed the structure-based *in silico* predictions. Compared with WT mKu, the mKuΔ12–15 mutant showed little to no depletion of the free DNA band at equimolar concentrations (5 nM) with FAM-labeled DNA. Even at a 200-fold molar excess (1 µM), mKuΔ12–15 displayed only weak DNA binding, with minimal loss of free DNA and limited formation of DNA–protein complexes. Notably, the DNA-binding activity of mKuΔ12–15 was comparably lower than that of the mKu-Ala mutant, despite residues 12–15 or the β-hairpin1 motif as a whole not directly contacting the DNA.

Our cryo-EM structures (Figure 1) revealed that, unlike the eukaryotic Ku70/80 heterodimer, mKu can independently drive DNA synapse formation. While cryo-EM provided a static visualization of this process in a frozen state, we extended our analysis to a more native condition in solution. To cross-validate this unique ability of mKu, we complemented the cryo-EM data with Förster resonance energy transfer (FRET) and hydrogen–deuterium exchange mass spectrometry (HDX-MS). These solution-based assays assess the stability and dynamics of the synapse, aspects that structure determination alone may not fully resolve. In the FRET assay, DNA concentration was held constant while mKu was titrated to saturation (Figure 5.d). Progressive addition of mKu resulted in a gradual decrease in donor emission accompanied by a corresponding increase in acceptor signal, consistent with fluorophore proximity through DNA synapse formation. Notably, the synapse remained stable throughout the experiment at 25 °C, underscoring both the assembly and stability of the DNA synaptic complex.

The stability and kinetics of the synaptic complex were further assessed using HDX-MS over a 0–100 min time course. This analysis provided residue-level insights into dynamics at both the mKu–mKu and DNA–mKu interfaces. Regions identified in the cryo-EM structures as critical for DNA binding within the central pore, as well as for DNA synapsis (β1 and helices 3–4), exhibited sustained protection throughout 0.1–100 min, consistent with stable contacts at these interfaces (Figure 3.a,b). Within the central pore, residues stretch 1–6 and 53–63, previously observed in close contact with DNA, showed strong protection. Similarly, the β-hairpin (residues 11–31), helix 3 (residues 111–124), and helix 4 (residues 194–210), which together form the synaptic interface, also displayed reduced deuterium uptake, supporting their role in synapse stabilization. By contrast, helix 7 (residues 252–273) at the C-terminal end of mKu became more exposed in the DNA-bound state compared to apo mKu. This region has been implicated in direct interaction with LigD, and its increased solvent accessibility upon DNA binding may prime mKu for recruitment of LigD, thereby enabling a seamless transition through the subsequent steps of the NHEJ pathway. Uptake profiles for residues essential for synapsis are shown in Figure 3.f, and overall HDX-MS statistics are provided in Supplementary Table 6.

### Capturing the initial phases of Mycobacterial NHEJ repair

To examine mKu dynamics after DNA damage, *M. smegmatis* (*Ku⁻ LigD⁻*) cells complemented with inducible plasmids expressing mKu–GFP were imaged. Laser micro irradiation within a defined ROI (0.8 × 1 µm) produced the expected loss of Hoechst 33342 signal due to photobleaching and triggered rapid accumulation of mKu–GFP at the damaged foci (Figure 7.a). The mKu–GFP signal subsequently dissipated, possibly due to mKu sliding along DNA molecules (8). In contrast, control ROIs without laser damage displayed diffuse mKu–GFP distribution, comparable to pre-irradiation images. The effect of photobleaching was also observed in the area surrounding the ROI. Quantification revealed a sharp decrease in the fluorescence intensity corresponding to Hoechst 33342, coincident with a spike in mKu–GFP intensity at the irradiated ROI (Figure 7.b). These observations demonstrate that mKu rapidly responds to double-strand breaks and dynamically localises at sites of DNA damage, suggesting its potential as a live-cell marker for bacterial DSBs.

**Figure 7.**
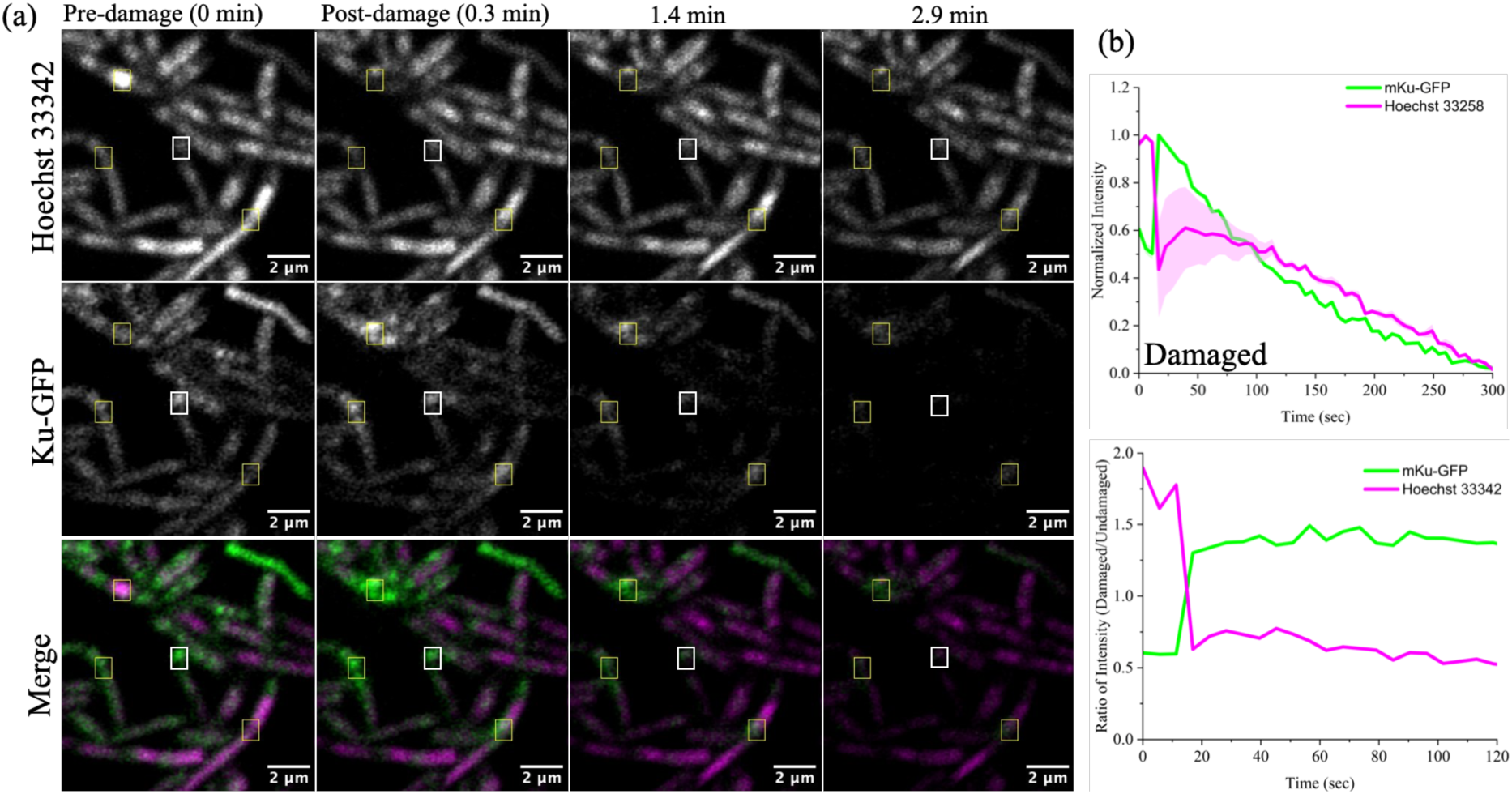
**| Live cell imaging captures DNA damage response of mKu** (a) Dynamics of mKu-GFP shown pre- and post-laser-induced DNA damage. The laser damage is focused within a 0.8 µm x 1 µm region (yellow box); the control ROI has the same dimensions (white box) but no laser damage. The dimensions of the boxes are scaled. DNA is stained with Hoechst 33342. (b) Quantification of fluorescence intensity fluctuation over time. Magenta and green traces represent the dynamics of Hoechst 33342 and mKu–GFP signals at the damaged ROI, respectively (top). The ratio of normalized intensities at damaged and undamaged ROI is plotted over time, with values above or below 1 indicating statistically significant changes (bottom).

## Discussion

In this study, we present the first cryo-electron microscopy (cryo-EM) structures of *Mycobacterium tuberculosis* Ku (mKu) bound to double-stranded DNA. These reconstructions provide direct visualization of the mKu–DNA complex and its higher-order oligomerization, revealing how mKu alone can orchestrate DNA synapsis in the absence of auxiliary repair factors (Figure 1). Our previous work showed that mKu forms filament-like assemblies on DNA (8), and the present structures now resolve the architecture of these assemblies, confirming that synapsis occurs through axial oligomerization along DNA. This bacterial mode of Ku-mediated end alignment contrasts sharply with the multi-protein synapsis seen in eukaryotic non-homologous end joining (NHEJ), where Ku70/80, DNA-PKcs, XRCC4, XLF, and Ligase IV act in concert to tether and process DNA ends (Figure 8) (11,28). By contrast, the *M. tuberculosis* two-component system, operating with mKu and Ligase D (LigD), illustrates a minimal yet robust bacterial NHEJ system.

**Figure 8.**
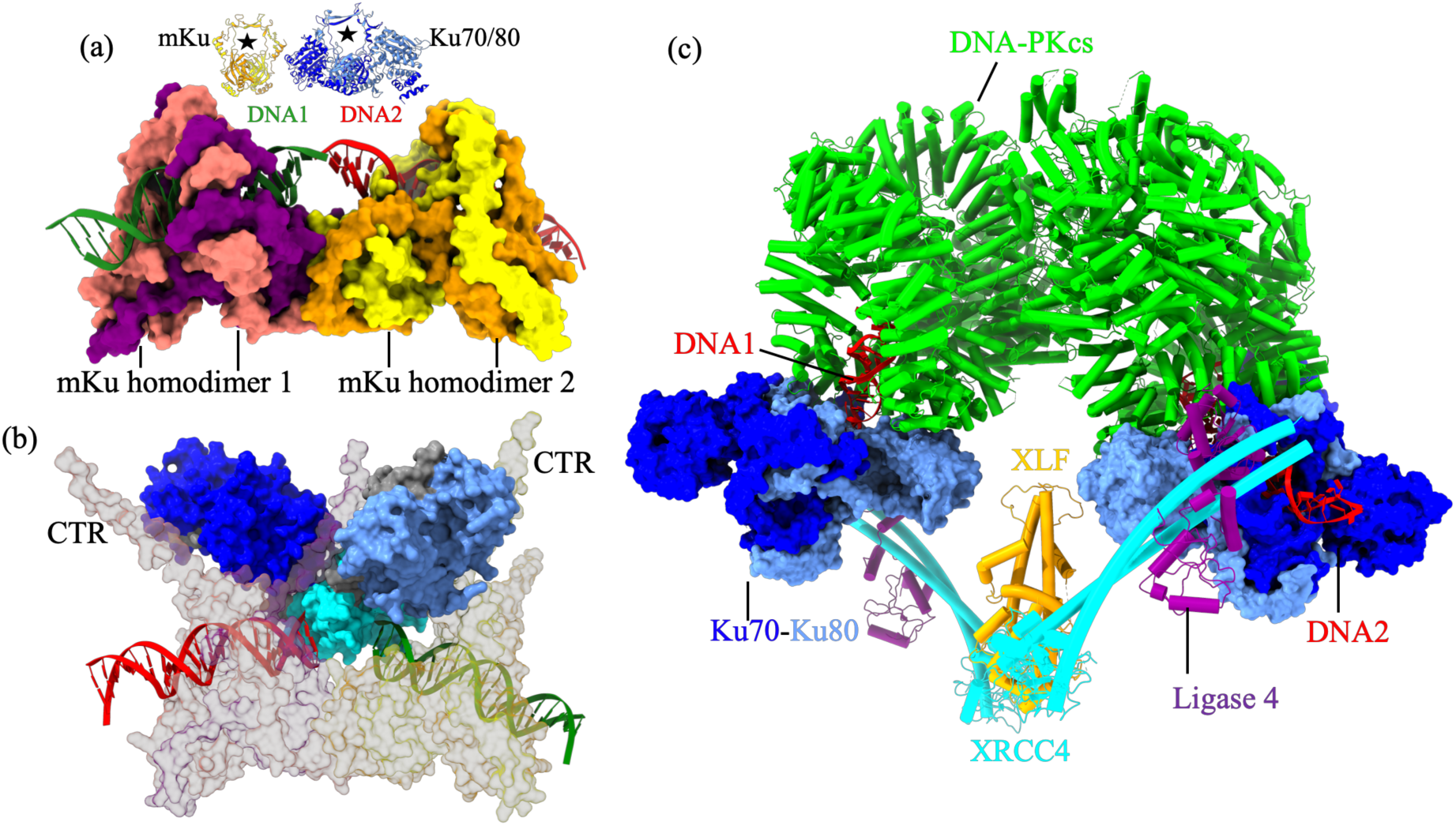
**| Relative simplicity of prokaryotic DNA synapsis** (a) Structural representation of the mKu-mediated DNA synaptic complex of *Mycobacterium tuberculosis*, showing mKu homodimer 1 (purple and magenta) and homodimer 2 (orange and yellow) binding DNA 1 (green) and DNA 2 (red). At the top the dimeric structures of mycobacterial and human Ku are compared to highlight the size differences in their respective complexes, with stars indicating the DNA-binding pores of each Ku dimer. (b) Modelled structure of the mycobacterial NHEJ complex (generated using the AlphaFold3 server), comprising mKu, DNA, and LigD. The three catalytic domains of LigD are depicted in cornflower blue (polymerase), cyan (phosphodiesterase), and dark blue (ligase). mKu dimers are shown in semi-transparent grey, and DNA molecules in red and green. The C-terminal region (CTR) of mKu directly contacts the ligase, consistent with experimental observations. (c) Structural representation of the human long-range DNA synaptic complex, with individual components colour-coded as indicated (PDB: 7NFC, Chaplin *et al.,* 2021).

Biochemical and biophysical analyses corroborate the structural model of mKu-mediated synapsis. FRET assays demonstrate persistent end-to-end proximity of two DNA termini in solution (Figure 5.d), while hydrogen–deuterium exchange mass spectrometry (HDX-MS) reveals sustained protection of both the DNA-binding pore and the synaptic β-hairpin 1 interface for over 100 minutes (Figure 3.a). Together, these findings indicate that the mKu-mediated synapse is remarkably stable and long-lived. This behaviour contrasts sharply with single-molecule studies on *Bacillus subtilis* Ku, where individual dimers mediate only transient (∼2 s) DNA bridging (12). The lack of structural information for *B. subtilis* Ku precludes a direct comparison with mKu; however, it suggests that the mycobacterial protein has evolved a distinct, cooperative mechanism to form a robust DNA alignment. As a single Ku dimer would be sterically constrained in accommodating two opposing DNA ends within its central pore, a continuous, multimeric arrangement is more plausible. This interpretation is reinforced by our cryo-EM maps, which reveal multiple adjacent dimers spanning the duplex to form a continuous synaptic filament. Collectively, these structural and biophysical data converge on a model in which mKu stabilizes DNA ends through cooperative filament formation rather than transient dimeric bridging.

Mechanistically, mKu first lodges at DNA termini before sliding inward, allowing successive dimers to assemble along the duplex (8). This progressive loading yields near-complete occupancy of DNA, forming a beads-on-a-string-like architecture visible in micrographs (Supplementary Figs. 3–4). Such continuous coating stabilizes DNA ends, promotes synapsis, and provides a scaffold for LigD recruitment. In *M. tuberculosis*, LigD acts as a multifunctional enzyme integrating three catalytic modules: an ATP-dependent ligase domain that seals nicks, a polymerase domain that can incorporate nucleotides, and a phosphoesterase (nuclease) domain that processes non-ligatable or damaged termini (29,30). Blunt-ended DNA substrates can be directly sealed by the ligase domain, whereas staggered or mismatched ends must first be trimmed to blunt ends by the nuclease. Once mKu aligns the termini, the accessibility of these LigD domains likely depends on its intrinsic interdomain flexibility, which allows dynamic repositioning around the synaptic scaffold. Our HDX-MS data show that DNA binding increases mKu’s C-terminal tail (CTR) flexibility, a region known to mediate LigD interaction (15,31), suggesting that DNA-induced conformational mobility facilitates transient fishing and capture of LigD at the synaptic junction (Figure 8.c). Together, these findings define a cooperative mechanism in which mKu provides the structural scaffold that aligns and stabilizes DNA ends, while LigD executes the catalytic steps of end processing and ligation. Thus, mKu functions as the rate-limiting architectural core of bacterial NHEJ, whereas LigD serves as its dynamic enzymatic component that completes the repair.

Structural analysis identifies two principal determinants that govern DNA binding and synapsis in mKu: (i) the positively charged central pore, which engages DNA in a sequence-independent manner, and (ii) the N-terminal β-hairpin 1 loop (residues 12–15), which mediates adjacent inter-dimer contacts. The central pore is lined with conserved Arg and Lys residues that electrostatically interact with the DNA sugar–phosphate backbone rather than bases, consistent with the canonical nut-and-bolt mode of DNA threading observed in other Ku homologs (Supplementary Figure 9). Although the overall topology of the Ku dimer, particularly the DNA-binding central pore, is conserved across species, the sequence homology is limited even among bacterial species (Supplementary Figure 10). Alanine substitution of key pore residues neutralizes the positive core of mKu, markedly impairing DNA binding (Figure 6) and corroborating a conserved charge-driven mechanism of DNA engagement.

In contrast, the β-hairpin 1 loop represents a distinctive structural element of mycobacterial Ku. Deletion of residues 12–15 shortens the loop, disrupts inter-dimer contacts (predicted by AlphaFold3), and abolishes both DNA binding and synapsis, as confirmed by comparative EMSA (Figure 6). Although the β-hairpin 1 does not directly contact DNA, its shortening markedly diminishes the DNA-binding affinity of mKu, underscoring how axial oligomerisation of mKu dimers impacts the stability of the DNA-protein supercomplex. This feature is absent in eukaryotic Ku (Figure 8.a,b), indicating a lineage-specific adaptation that enables robust DNA end bridging in *M. tuberculosis*. Thus, unlike eukaryotic homologues, mKu’s DNA-binding affinity arises cumulatively from direct electrostatic engagement with DNA and, more importantly, from cooperative stabilization at the synaptic interface, where disruption of the latter compromises both DNA association and end bridging.

We further observe that successive mKu dimers are not oriented perpendicularly to the DNA axis; rather, each additional dimer induces a helical rotation of ∼137° on linear dsDNA and ∼127° on shDNA. This gradual rotation reduces steric hindrance between adjacent dimers and opposite DNA ends, thereby promoting a compact and continuous synaptic organization. The synaptic interface thus acts as a flexible hinge, permitting controlled rotation around the DNA axis to minimize steric clashes and optimize packing density. Such behaviour resonates with intrinsic DNA helical geometry, where twist and base stacking govern conformational stability. An open question remains whether the loop architecture of β-hairpin 1 itself or its constituent residues is the principal determinant of this twist modulation. AlphaFold3-generated models with alanine substitutions at residues 12–15 retained a predicted synaptic arrangement (Supplementary Figure 11), suggesting that the loop’s topology, rather than specific side-chain chemistry, may dictate the helical orientation. Experimental validation using the mKu Ala (12–15) construct will be required to confirm this structural prediction.

In solution, mKu exists as a stable homodimer that oligomerizes only upon engaging linear DNA. The dimer adopts a clasped human hand-like architecture, stabilized by β-hairpin and helix–turn–helix motifs, which compact the core. In contrast, the flexible C-terminal region (CTR; residues 228–273) lacks defined density in cryo-EM maps, consistent with its elevated deuterium uptake, which supports its role as a dynamic tether for LigD. Future structural studies on the complete mKu–DNA–LigD complex will be required to visualize this interaction and elucidate how LigD domains reorganize around the mKu scaffold during repair.

Live-cell confocal imaging of *Mycobacterium smegmatis* expressing mKu–GFP revealed rapid accumulation of mKu at laser-induced double-strand breaks (Figure 7), consistent with its high affinity for linear DNA ends. Over time, the redistribution of mKu–GFP fluorescence away from the break site suggests lateral diffusion along the DNA duplex, a mechanism that remains to be directly validated by single-molecule fluorescence imaging. Given the large size of the *M. tuberculosis* genome (≈4.4 Mbp), the cellular abundance of mKu may be insufficient to span entire genomic tracts, raising the possibility that mKu availability could constitute a rate-limiting parameter for repair. Future studies employing single-molecule tracking in live cells could define the stoichiometry, diffusion behaviour, and turnover dynamics of mKu during DNA repair.

Despite providing high-resolution insights into prokaryotic Ku–DNA synaptic assemblies, certain limitations remain. Local conformational heterogeneity within the oligomers limited resolution at the DNA termini and in the CTR, precluding direct visualization of the LigD-recruitment interface. Moreover, while deletion and alanine-substitution mutants validate the structural and functional roles of the central pore and β-hairpin 1 in DNA binding and synapsis, future assays coupling LigD-dependent ligation with these mutants will be essential to link specific structural perturbations to NHEJ efficiency. The integration of cryo-EM with in situ single-molecule imaging could further elucidate how mKu dynamics are regulated in a cellular context.

In summary, this work defines a bacterial-specific mechanism of Ku-mediated DNA synapsis. Through cooperative oligomerization along DNA, mKu autonomously aligns DNA ends without the auxiliary factors required in eukaryotic NHEJ. The β-hairpin 1 loop, comprising only four residues (12–15), emerges as a critical determinant of synapsis and stability. This structural adaptation, absent in human Ku70/80, highlights an evolutionary divergence that could be leveraged to selectively probe bacterial DNA repair without cross-reactivity in human cells. By integrating structural, biochemical, and cellular analyses, our study provides a mechanistic framework for understanding bacterial NHEJ and sets the stage for exploring DNA-repair vulnerabilities in persistent *M. tuberculosis* infections.

## Data availability

For ds-DNA and sh-DNA, the cryo-EM density maps have been deposited in the Electron Microscopy Data Bank (EMDB) under IDs EMD- 43184, -43186, and EMD- 42978, -43185, respectively. Coordinates are deposited in the Protein Data Bank (PDB) under IDs PDB-8VF5, 8VF2, 8V53, and 8VF4, respectively. The HDX-MS data are available from ProteomeXchange via the PRIDE partner repository with the identifier PXD060776.

## Author contributions

J.B. conceived the study and performed biochemical and biophysical experiments, light microscopy, and cryo-EM sample preparation, data processing, single-particle reconstruction, and model building under the supervision of E.H., A.K.D. and I.R. C.S.A. acquired and supervised HDX-MS data analysis performed by J.B. K.S. and J.B. acquired light microscopy data, which J.B. processed and analyzed under the guidance of P.J.M. and E.H. A.S. purified mKu mutant constructs and performed EMSA under J.B.’s supervision. J.B. and I.R. wrote the manuscript with input from all authors.

## Acknowledgments

We sincerely thank the University of Melbourne’s research platforms, including the Ian Holmes Imaging Centre (IHC) for access to the Cryo-EM platform, the Biological Optical Microscopy Platform (BOMP) for light microscopy facilities, the Mass Spectrometry and Proteomics Facility for HDX-MS data collection, and the Melbourne Protein Characterisation (MPC) Facility for support with protein purification and characterization. We are especially grateful to Dr. Hamish Brown and Dr. Sepideh Valimehr (IHC) for their assistance with electron microscopy training and data collection and to Dr. Mohammad Hossein Tanipour for his guidance on EM grid preparation. We also acknowledge Martin Fleming for technical support and Sean Crosby for supercomputing assistance. Data storage and processing were conducted in the Rouiller lab and on the Spartan supercomputer hosted by Melbourne Research Computing. We thank Prof. Aidan Doherty (University of Sussex) for providing the mKu-containing recombinant plasmid (pET16b-mKu) and Prof. Michael S. Glickman (Memorial Sloan Kettering Cancer Center, New York) for the *Mycobacterium smegmatis* mc² 155 Δku ΔligD strain (Msm 153). We acknowledge the Melbourne India Postgraduate Academy (MIPA) program for facilitating the joint collaboration and doctoral degree program (granted to JB) between IIT Kharagpur and the University of Melbourne.

