## Supplementary data for "Bringing the Ends together: Cryo-EM structures of mycobacterial Ku in complex with DNA define its role in NHEJ synapsis"

### Supplementary Figures:

### Supplementary Figure 1 | Sequencing confirmation of mKu, mKu Ala, mKu 12-15

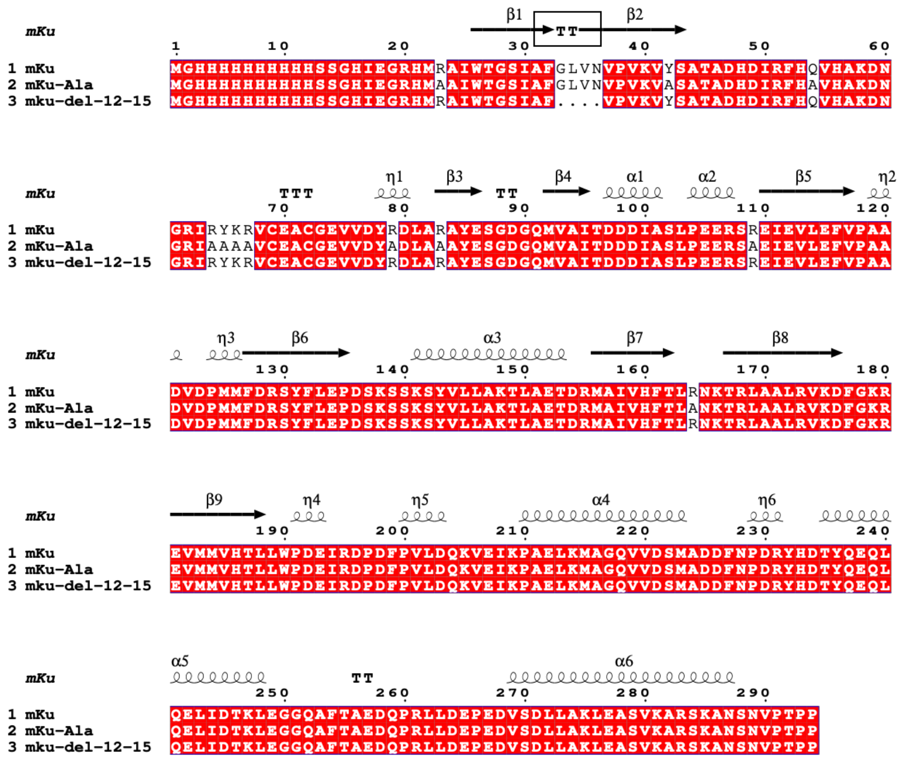

**Supplementary Figure 1 | Sequencing confirmation of mKu and its mutants:** A topology diagram is shown at the top, aligned with the sequence alignment below. Residues 12–15, which form the loop region of β-hairpin-1, are highlighted with a black box. Eleven residues within the DNA-binding core were substituted with alanine in the DNA-binding–impaired mutant (mKu-Ala).

### Supplementary Figure 2 | Purification of mKu and its mutants

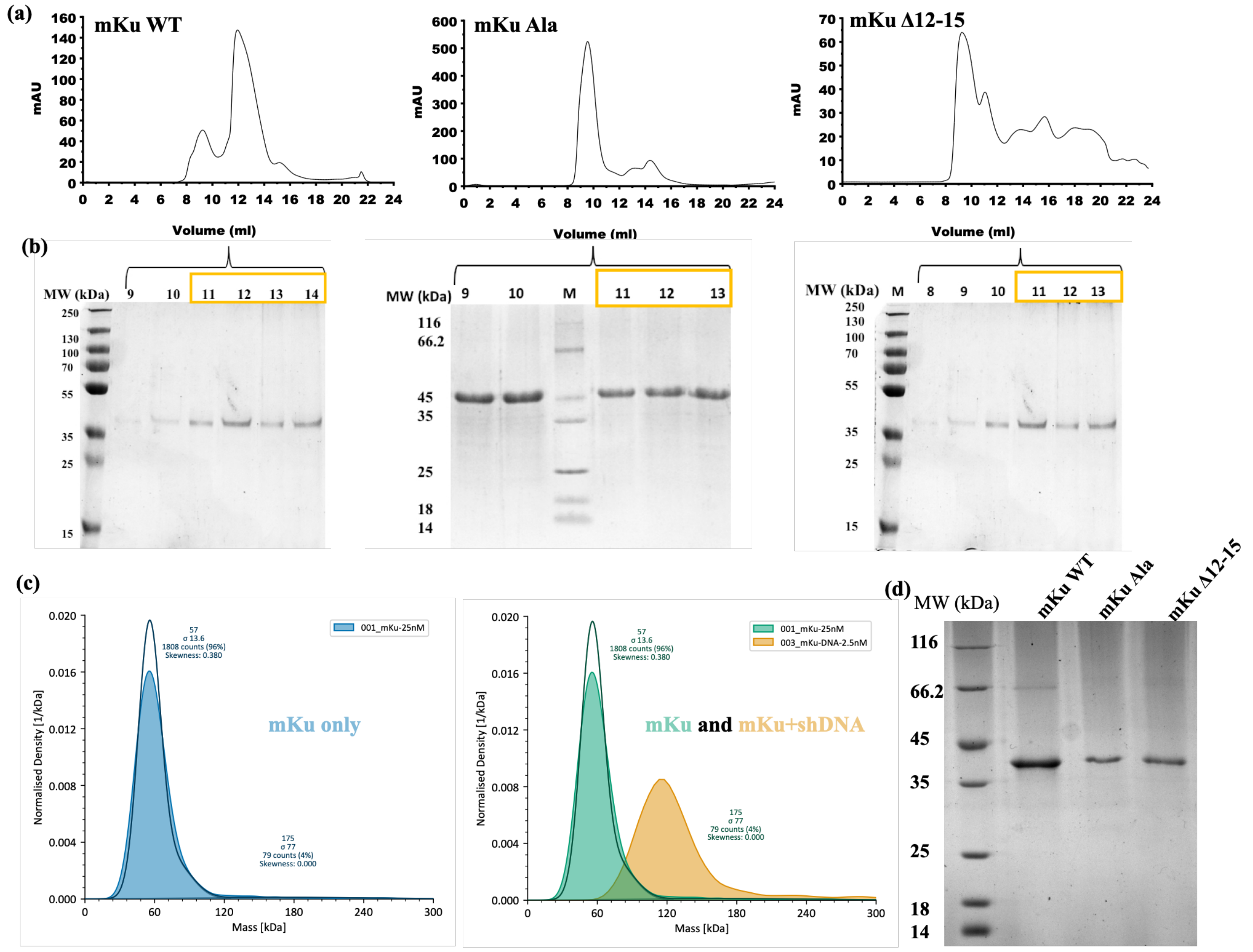


**Supplementary Figure 2 | Purification of mKu and its mutants**: (a) Analytical size-exclusion chromatography (SEC) profiles of mKu WT, mKu Ala, and mKu Δ12–15 proteins on a Superdex-200 10/300 GL column (Cytiva). Each protein elutes in two phases: an early peak at 8–10 ml with $OD_{260/280} > 1$, and a later peak at 11–14 ml with $OD_{260/280} \leq 0.6$. Both the fractions (early and late) show a single band in SDS PAGE. The early-eluting fractions correspond to higher-order oligomers of mKu bound to endogenous *E. coli* DNA, reflecting the high DNA-binding affinity of mKu. For experimental assays, fractions from 11–14 ml were pooled due to minimal DNA contamination. (b) SDS-PAGE analysis of SEC fractions (9–14 ml) for mKu WT and mutants, confirming protein purity in the pooled fractions. (c) Mass photometry analysis of mKu alone and mKu incubated with short DNA. The protein-only sample shows a molecular weight of ~60 kDa (blue), consistent with the theoretical molecular weight of 66.8 kDa. Upon addition of DNA, the population shifts to ~175 kDa (yellow), indicating formation of mKu–DNA oligomers. The broad peak distribution (mKu+DNA) reflects the heterogeneity of the sample due to higher-order oligomerisation. (d) SDS-PAGE analysis of purified mKu WT and mutant proteins after sequential purification, showing the final samples used for experiments. A faint band at ~66 kDa corresponds to mKu dimers, a characteristic feature of mKu proteins arising from their partial resistance to SDS and high thermal stability(8).

### Supplementary Figure 3 | Comparison between mKu-dsDNA and mKu-shDNA complexes

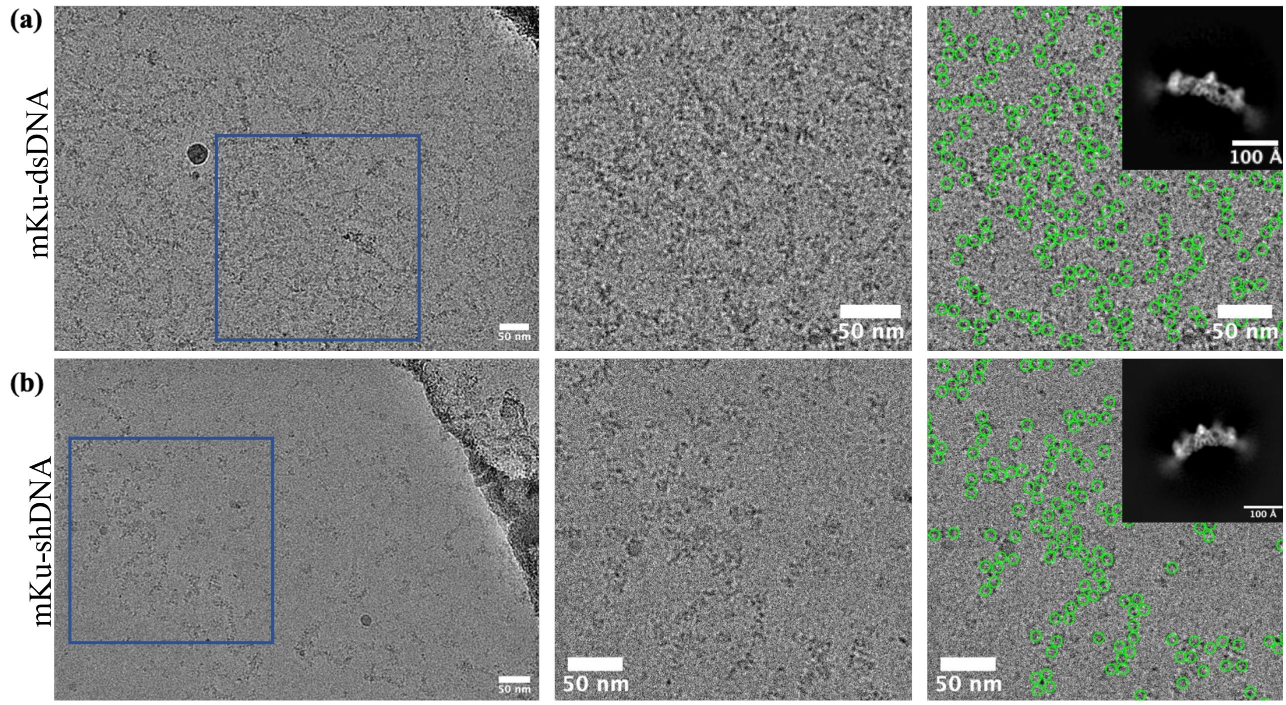


**Supplementary Figure 3 | Comparison between cryo-EM datasets of mKu complex:** Analysis of the mKu-dsDNA (a) and mKu-shDNA (b) oligomers. The first panel displays the raw cryo-EM micrographs for each complex. The subsequent panels show an enlarged section of the micrographs highlighting the particles, followed by the same section with particle picks marked with green circles (100 Å diameter) to estimate oligomer size. On the rightmost panel, representative 2D class averages illustrate the differences in bending or curvature between the mKu-dsDNA and mKu-shDNA complexes.

### Supplementary Figure 4 | Cryo-EM data processing workflow

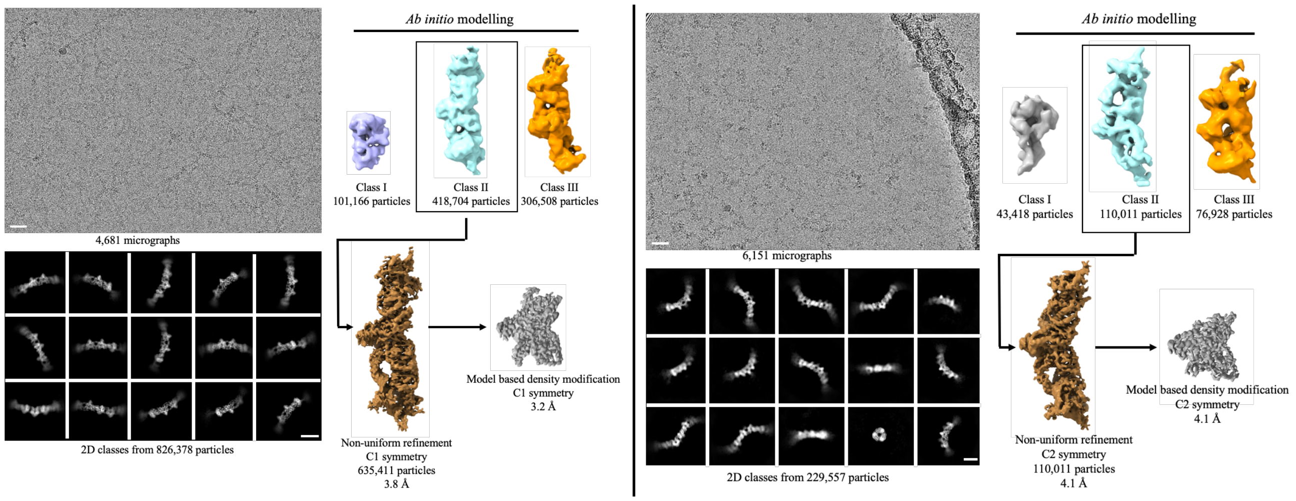


**Supplementary Figure 4 | Cryo-EM data processing workflow for mKu-DNA complexes:** The left and right panels illustrate the data processing workflows for the mKu-dsDNA and mKu-shDNA datasets, respectively. Each workflow sequentially presents the raw micrographs, 2D class averages, ab initio models, and refined models, showcasing the progression from raw data to high-resolution structures.

### Supplementary Figure 5 | Map validation and resolution estimation

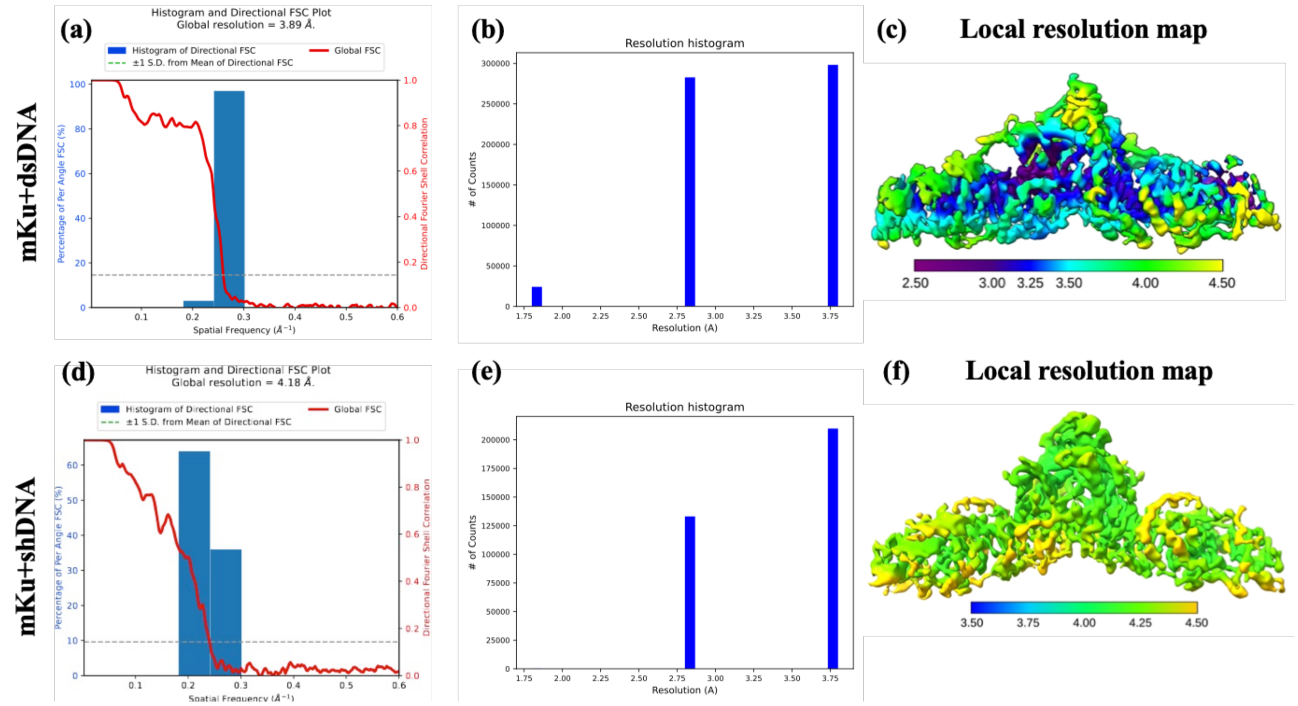


**Supplementary Figure 5 | Map validation and resolution estimation for mKu-DNA complexes:** (a & d) 3D Fourier Shell Correlation (FSC) plot from CryoSparc showing the global resolution of the map; (b & e) resolution histogram generated by ResMap; (c & f) local resolution map calculated using Phenix, indicating the range of resolution values across the respective maps.

### Supplementary Figure 6 | FRET showing mKu-mediated DNA synapsis

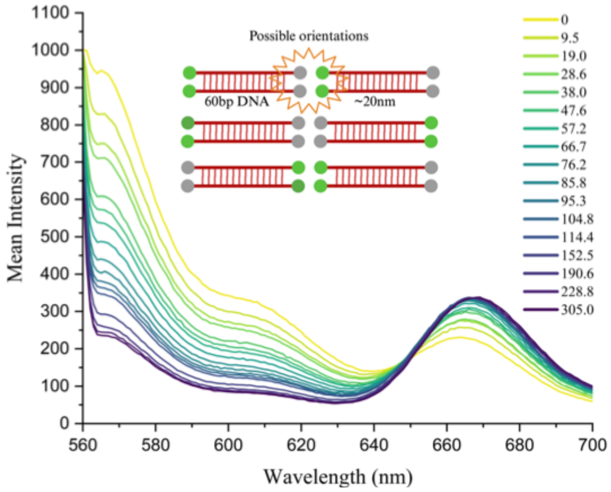


**Supplementary Figure 6 | FRET analysis of DNA substrate interactions in the presence of mKu:** Wavelength vs. fluorescence intensity plot illustrating Förster Resonance Energy Transfer (FRET) between DNA substrates labelled with donor (Cy3) and acceptor (Cy5) fluorophores. The DNA concentration was fixed at 250 µM, while mKu was titrated incrementally from 0 to 305 µM. The schematic representation depicts the synaptic orientations enabling successful FRET, with the donor fluorophore shown in green and the acceptor in grey.

### Supplementary Figure 7 | Live cell imaging captures post-DNA damage response of mKu

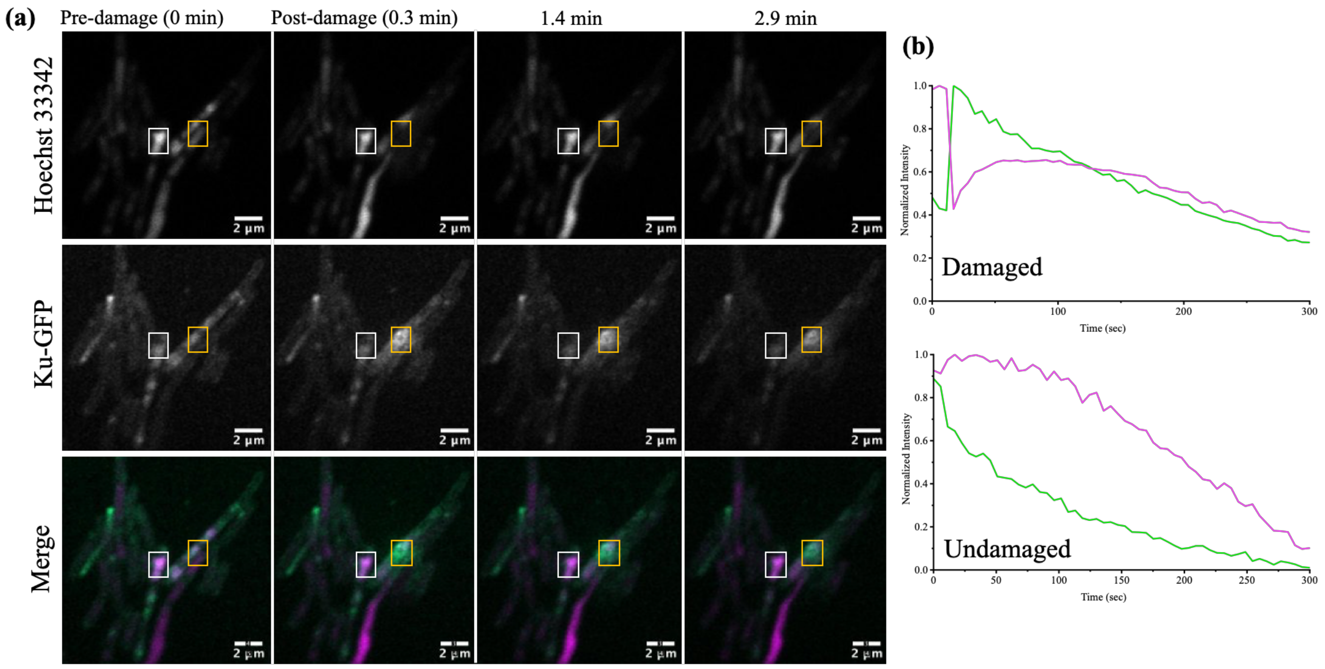


**Supplementary Figure 7 | Live cell imaging captures DNA damage response of mKu**: (a) Dynamics of mKu-GFP shown pre- and post-laser-induced DNA damage. The laser damage is focused within a 0.8 µm x 1 µm region (yellow box); the control ROI has the same dimensions (white) but no laser damage. The dimensions of the boxes are not scaled. DNA is stained with Hoechst 33342. (b) Intensity quantification over time, with magenta and green lines representing the dynamics of DNA and mKu-GFP signals at damaged and undamaged ROI, respectively.

### Supplementary Figure 8 | mKu-mKu dimer interface

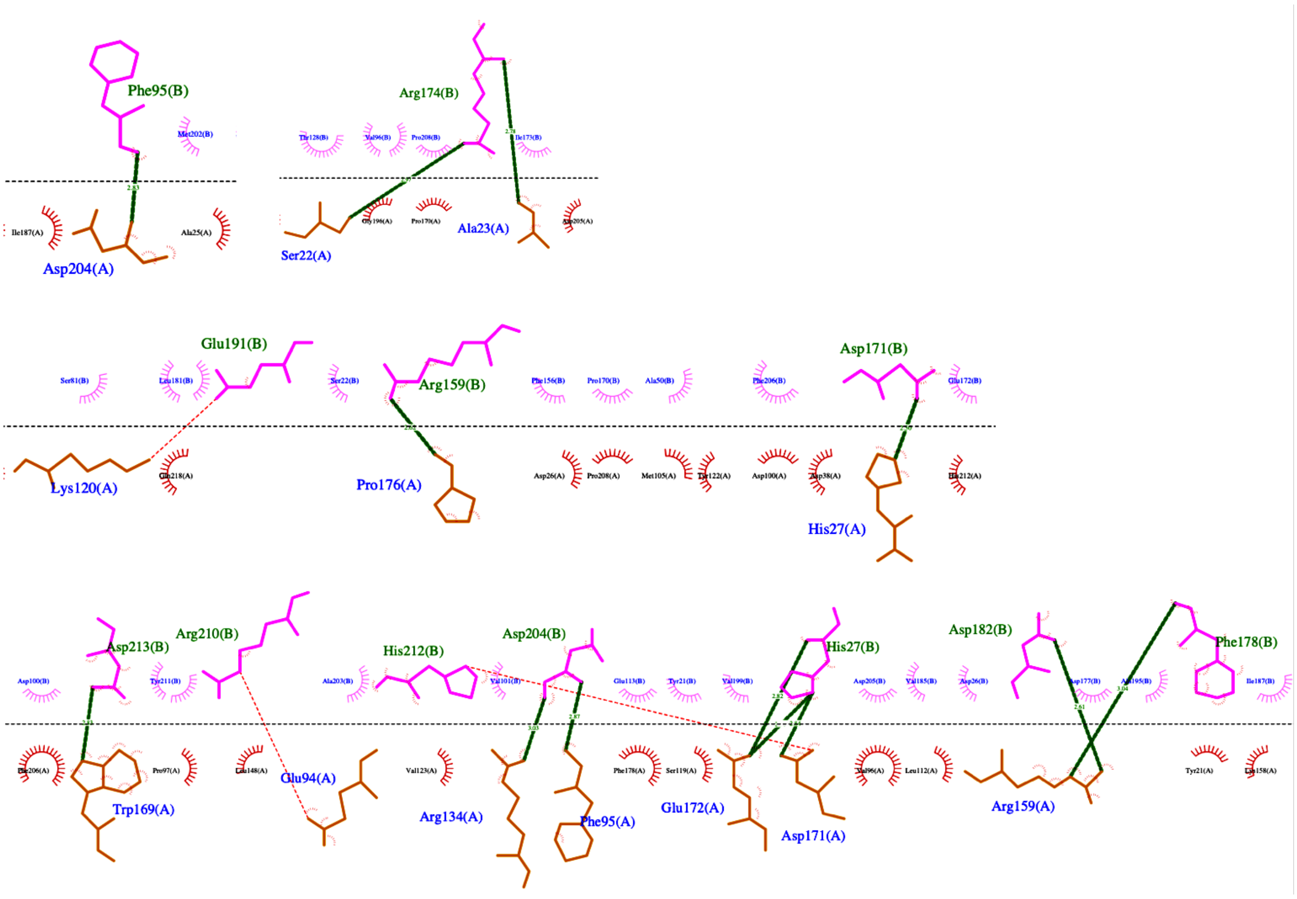


**Supplementary Figure 8 | Protein-protein dimerization interface of the mKu dimer:** The dimerization interface is depicted using a 2D contact map generated with LigPlot+. Amino acid residues from the two monomeric chains are coloured purple and brown, respectively. Neighbouring hydrophobic residues are represented as spikes. Hydrogen bonds are shown as green dashed lines, while salt bridges are indicated with red dashed lines.

### Supplementary Figure 9 | The Core DNA-binding domain of Ku is structurally conserved

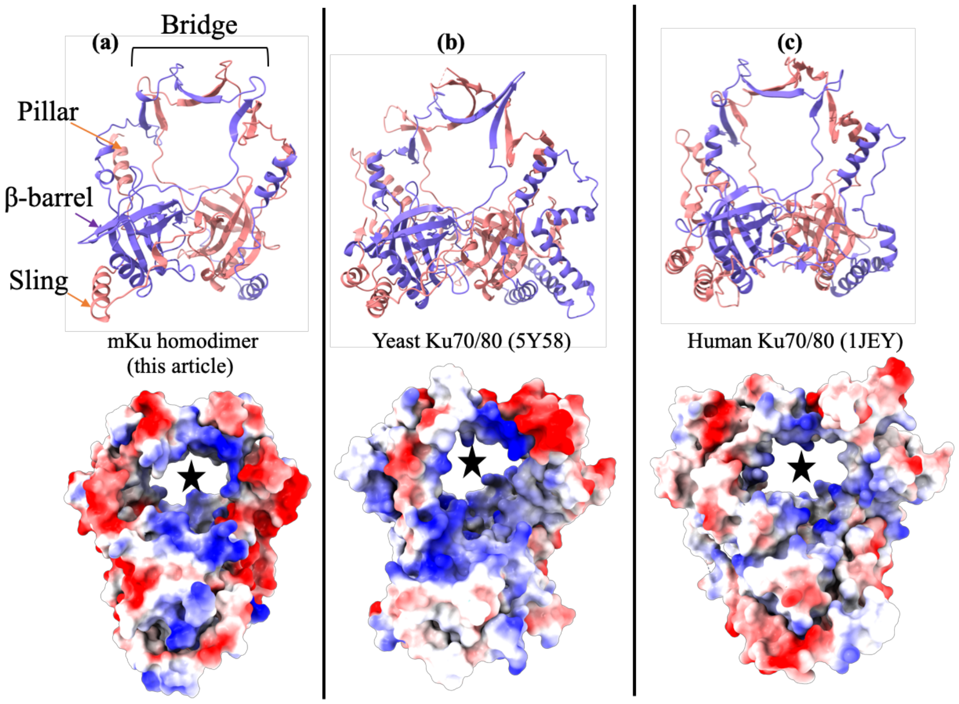


**Supplementary Figure 9 | Comparison of the core DNA-binding domain of Ku proteins:** Experimentally determined structures reveal the similarity of the DNA-binding domain across mKu (PDB: 8VF5; this study) (a), yeast Ku70/80 (PDB: 5Y58) (b), and human Ku70/80 (PDB: 1JEY) (c). Additional domains in yeast and human Ku were removed for clarity. The bottom panels illustrate the electrostatic charge distributions, with positive to negative charges represented by red to blue gradients. The DNA-binding pores, lined with positively charged residues, are marked with a star symbol.

### Supplementary Figure 10 | Comparison of DNA binding residues of Ku (Mtb, Human and Yeast)

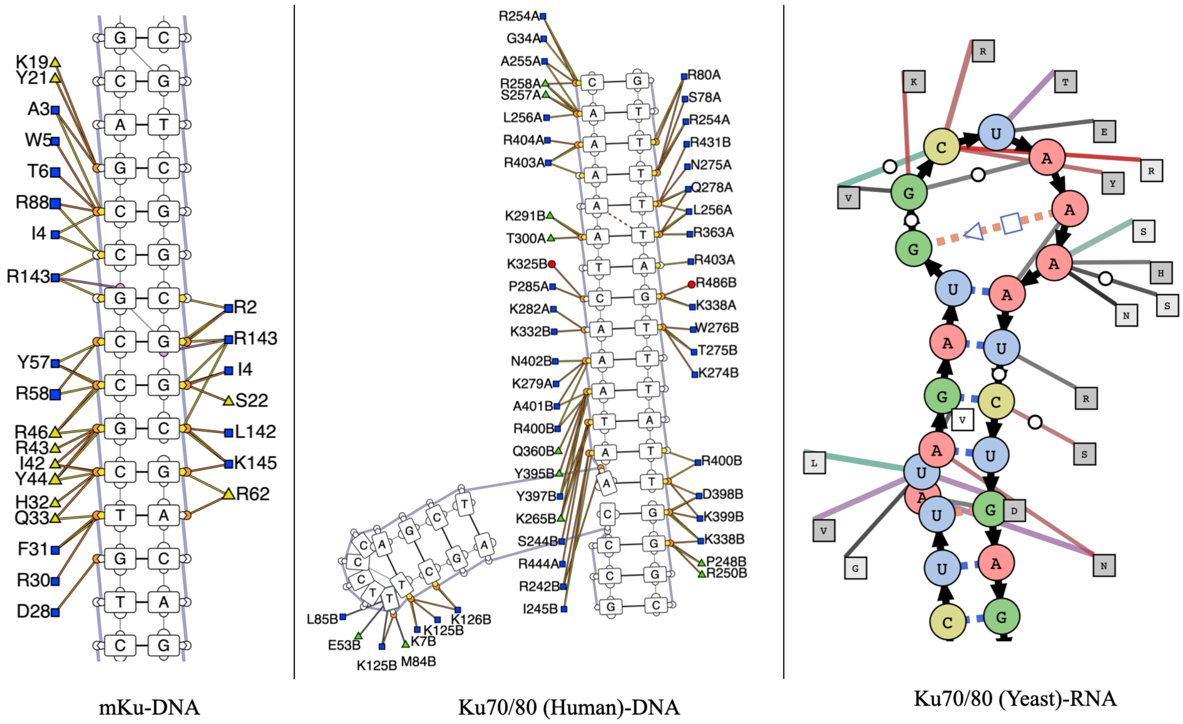


**Supplementary Figure 10 | Comparison of the DNA-binding residues of Ku proteins:** Experimentally determined structures highlight the DNA-binding residues of mKu (PDB: 8VF5; this study), human Ku70/80 (PDB: 1JEY) and yeast

Ku70/80 (PDB: 5Y58). The yeast Ku70/80 structure is shown in complex with RNA. Despite the absence of sequence homology at the DNA/RNA-binding pore, the binding mode is conserved, with most contacts mediated by positively charged residues (Arg and Lys) interacting with the sugar–phosphate backbone.

### Supplementary Figure 11 | Predictive structural modelling of mKu Ala (12-15) DNA synaptic complex

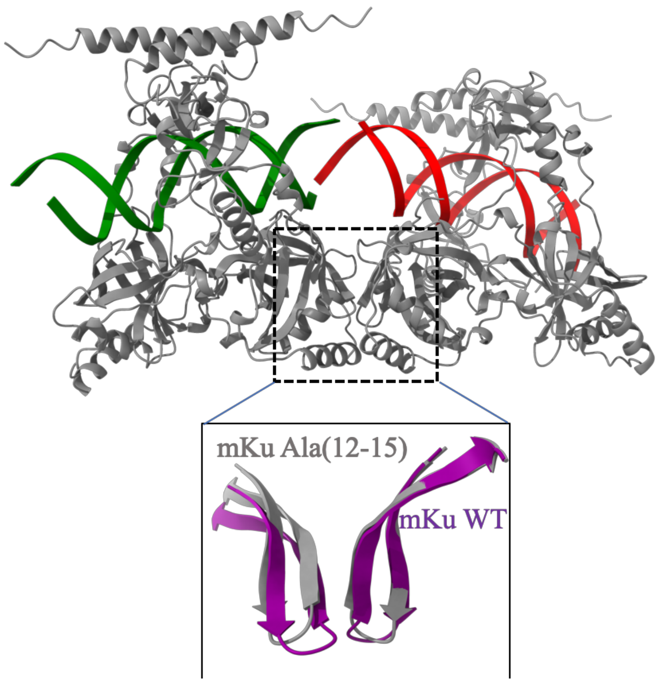


**Supplementary Figure 11 | Predictive structural modelling of the mKu Ala (12–15) DNA synaptic complex:** AlphaFold3-predicted model of the mKu Ala (12–15) mutant bound to two dsDNA duplexes. The synaptic interface remains intact despite alanine substitution, in contrast to the Δ (12–15) mutant, where synapsis is disrupted. This suggests that the overall β-hairpin 1 topology, rather than specific side-chain interactions, is the primary determinant of synapsis. The β-hairpin 1 of WT (purple) and Ala (12–15) (grey) mKu are superimposed to illustrate deviations from the wild-type synaptic conformation.

### Supplementary Tables:

### Supplementary Table 1 | Composition of buffers used in this study

| Buffer | Composition |
|---|---|
| Resuspension buffer | 50 mM Tris pH 7.5, 250 mM NaCl, 10% sucrose, PMSF 17 $\mu g \cdot ml^{-1}$, benzamidine 34 $\mu g \cdot ml^{-1}$ |
| IMAC wash buffer-1 | 50 mM Tris pH 7.5, 400 mM NaCl, 10 mM imidazole, 10% glycerol, PMSF 17 $\mu g \cdot ml^{-1}$, benzamidine 34 $\mu g \cdot ml^{-1}$ |
| IMAC wash buffer-2 | 50 mM Tris pH 7.5, 400 mM NaCl, 45 mM imidazole, 10% glycerol, PMSF 17 $\mu g \cdot ml^{-1}$, benzamidine 34 $\mu g \cdot ml^{-1}$ |
| IMAC elution buffer | 50 mM Tris pH 7.5, 400 mM NaCl, 300 mM imidazole, 10% glycerol, PMSF 17 $\mu g \cdot ml^{-1}$, benzamidine 34 $\mu g \cdot ml^{-1}$ |
| IEX buffer-1 | 50 mM Tris pH 7.5, 50 mM NaCl, 2 mM DTT, 10% glycerol, PMSF 17 $\mu g \cdot ml^{-1}$, benzamidine 34 $\mu g \cdot ml^{-1}$ |
| IEX buffer-2 | 50 mM Tris pH 7.5, 1 M NaCl, 2 mM DTT, 10% glycerol, PMSF 17 $\mu g \cdot ml^{-1}$, benzamidine 34 $\mu g \cdot ml^{-1}$ |
| SEC buffer | 25 mM HEPES pH 7.5, 150 mM NaCl, 2 mM DTT, 10% glycerol, PMSF 17 $\mu g \cdot ml^{-1}$, benzamidine 34 $\mu g \cdot ml^{-1}$ |

### Supplementary Table 2 | Cryo-EM data collection, refinement, and validation statistics

| | mKu-linear DNA | | mKu-hairpin DNA | |
|---|---|---|---|---|
| | Unit complex | Supercomplex | Unit complex | Supercomplex |
| **Data collection and processing** | PDB: 8VF5<br>EMD: 43186 | PDB: 8VF2<br>EMD: 43184 | PDB: 8V53<br>EMD: 42978 | PDB: 8VF4<br>EMD: 43185 |
| Micrographs | 6151 | | 4682 | |
| Magnification | 105,000 | | 105,000 | |
| Voltage (kV) | 300 | | 300 | |
| Electron exposure ($e^{-}$/Å$^2$) | 40 | | 40 | |
| Defocus range (μm) | -1.5 to -2.5 | | -1.5 to -2.5 | |
| Pixel size (Å) | 0.833 | | 0.833 | |
| Symmetry imposed | C1 | | C2 | |
| Initial particle (no.) | ~ 4,800,000 | | ~ 1,150,000 | |
| Final particle (no.) | 635,411 | | 110,011 | |
| Map resolution (Å) | 3.7 | 3.7 | 4.1 | 4.1 |
| FSC threshold | 0.143 | 0.143 | 0.143 | 0.143 |
| Map resolution range (Å) | 2.0-3.9 | 2.0 - 5.7 | 4.0 - 5.2 | 4.0 - 5.2 |
| **Refinement** | | | | |
| Model resolution (Å) | 3.7 | 3.7 | 4.1 | 4.1 |
| FSC threshold | 0.143 | 0.143 | 0.143 | 0.143 |
| Map sharpening B factor (Å$^2$) | 168.6 | 168.6 | 161.5 | 161.5 |
| **Model composition** | | | | |
| Non-hydrogen atoms | 4513 | 13,367 | 4281 | 12,777 |
| Protein residues | 453 | 1359 | 453 | 1359 |
| DNA | 42 | 118 | 31 | 90 |
| **B factors (Å$^2$)** | | | | |
| Protein | 65.04 | 84.86 | 150.25 | 143.92 |
| DNA | 75.53 | 153.25 | 139.88 | 216.43 |
| **R.M.S.D.** | | | | |
| Bond lengths (Å) | 0.004 | 0.004 | 0.004 | 0.004 |
| Bond angles (°) | 0.91 | 0.89 | 0.92 | 0.89 |
| **Validation** | | | | |
| MolProbity score | 1.95 | 1.9 | 1.96 | 1.99 |
| Clash score | 10.39 | 11.48 | 14.58 | 16.72 |
| Rotamer outliers (%) | 0.25 | 0.34 | 0.76 | 0.42 |
| **Ramachandran plot** | | | | |
| Favoured (%) | 93.76 | 95.4 | 95.77 | 96.14 |
| Allowed (%) | 6.24 | 4.6 | 4.23 | 3.86 |
| Disallowed (%) | 0 | 0 | 0 | 0 |

### Supplementary Table 3 | List of interacting residues for mKu-mKu dimer

| Atom no. | Atom name | Res name | Res no. | Chain ID | | Atom no. | Atom name | Res name | Res no. | Chain ID | Distance (Å) |
|---|---|---|---|---|---|---|---|---|---|---|---|
| 302 | N | Ala | 61 | B | --- | 169 | O | Val | 34 | A | 2.194 |
| 355 | N | Arg | 41 | A | --- | 187 | OE2 | Glu | 49 | B | 2.334 |
| 126 | N | His | 32 | A | --- | 211 | O | Ala | 63 | B | 2.476 |
| 326 | OH | Tyr | 211 | A | --- | 266 | OE1 | Glu | 94 | B | 2.477 |
| 135 | NE2 | His | 32 | A | --- | 385 | OE2 | Glu | 65 | B | 2.496 |
| 443 | NE2 | His | 27 | A | --- | 556 | OD2 | Asp | 171 | B | 2.496 |
| 455 | NH2 | Arg | 210 | A | --- | 267 | OE2 | Glu | 94 | B | 2.541 |
| 713 | NZ | Lys | 120 | B | --- | 722 | OE2 | Glu | 191 | A | 2.571 |
| 115 | NH2 | Arg | 159 | A | --- | 393 | OD2 | Asp | 182 | B | 2.61 |
| 589 | NH1 | Arg | 159 | B | --- | 477 | O | Pro | 176 | A | 2.622 |
| 454 | NH1 | Arg | 210 | A | --- | 267 | OE2 | Glu | 94 | B | 2.627 |
| 74 | N | Glu | 49 | A | --- | 147 | O | Arg | 41 | B | 2.653 |
| 83 | N | Val | 47 | B | --- | 200 | O | Arg | 43 | A | 2.725 |
| 154 | NH2 | Arg | 41 | B | --- | 195 | OE1 | Glu | 65 | A | 2.755 |
| 122 | ND1 | His | 27 | B | --- | 104 | OE2 | Glu | 172 | A | 2.757 |
| 213 | N | Val | 34 | B | --- | 248 | O | Ala | 61 | A | 2.766 |
| 234 | N | Arg | 174 | B | --- | 497 | O | Ala | 23 | A | 2.781 |
| 197 | N | Arg | 43 | A | --- | 86 | O | Val | 47 | B | 2.791 |
| 279 | N | Ala | 63 | A | --- | 67 | O | His | 32 | B | 2.8 |
| 116 | N | His | 27 | B | --- | 104 | OE2 | Glu | 172 | A | 2.817 |
| 466 | N | Asp | 204 | A | --- | 531 | O | Phe | 95 | B | 2.826 |
| 23 | NE1 | Trp | 169 | A | --- | 143 | OD2 | Asp | 213 | B | 2.833 |
| 122 | ND1 | His | 27 | B | --- | 353 | OD1 | Asp | 171 | A | 2.842 |
| 208 | N | Ala | 63 | B | --- | 129 | O | His | 32 | A | 2.866 |
| 307 | N | Asp | 204 | B | --- | 602 | O | Phe | 95 | A | 2.867 |
| 64 | N | His | 32 | B | --- | 282 | O | Ala | 63 | A | 2.867 |
| 245 | N | Ala | 61 | A | --- | 216 | O | Val | 34 | B | 2.891 |
| 38 | NH1 | Arg | 43 | B | --- | 54 | O | Val | 47 | A | 2.894 |
| 244 | NH2 | Arg | 174 | B | --- | 1055 | OG | Ser | 22 | A | 2.973 |
| 51 | N | Val | 47 | A | --- | 32 | O | Arg | 43 | B | 3.029 |
| 704 | NH2 | Arg | 134 | A | --- | 310 | O | Asp | 204 | B | 3.034 |
| 112 | NE | Arg | 159 | A | --- | 43 | O | Phe | 178 | B | 3.037 |
| 29 | N | Arg | 43 | B | --- | 54 | O | Val | 47 | A | 3.059 |
| 508 | NE2 | His | 35 | A | --- | 421 | SG | Cys | 51 | B | 3.062 |
| 9 | NE1 | Trp | 169 | B | --- | 233 | OD2 | Asp | 213 | A | 3.069 |
| 618 | NZ | Lys | 37 | A | --- | 802 | OE2 | Glu | 53 | B | 3.105 |
| 166 | N | Val | 34 | A | --- | 305 | O | Ala | 61 | B | 3.213 |
| 275 | NE | Arg | 46 | B | --- | 405 | OH | Tyr | 44 | A | 3.306 |

**Supplementary Table 4 | Impact of mutation of the critical residues in the DNA-protein interface**

| Chain | Wild Residue | Residue Position | Mutant Residue | RSA(%) | Predicted ΔΔG | Outcome |
|---|---|---|---|---|---|---|
| A | Y | 21 | A | 11.2 | -0.115 | Destabilizing |
| A | Q | 33 | A | 16.4 | -2.616 | Highly destabilizing |
| A | R | 43 | A | 37.7 | -2.432 | Highly destabilizing |
| A | Y | 44 | A | 14.3 | -2.665 | Highly destabilizing |
| A | R | 46 | A | 12.1 | -4.149 | Highly destabilizing |
| A | R | 58 | A | 77.5 | -1.826 | Destabilizing |
| A | R | 62 | A | 32.1 | -2.702 | Highly destabilizing |
| A | R | 66 | A | 44.9 | -0.869 | Destabilizing |
| B | Y | 21 | A | 11.2 | -0.322 | Destabilizing |
| B | Q | 33 | A | 16.4 | ·2.258 | Highly Destabilizing |
| B | R | 43 | A | 37.7 | -3.498 | Highly Destabilizing |
| B | Y | 44 | A | 14.3 | -3.427 | Highly Destabilizing |
| B | R | 46 | A | 12.1 | -3.338 | Highly Destabilizing |
| B | R | 58 | A | 77.5 | -1.566 | Destabilizing |
| B | R | 62 | A | 32.1 | -3.046 | Highly Destabilizing |
| B | R | 88 | A | 44.9 | -0.934 | Destabilizing |

**Supplementary Table 5 | Impact of mutation of the critical residues in the synaptic interface**

| Chain | Wild Residue | Residue Position | Mutant Residue | RSA(%) | Predicted ΔΔG | Outcome |
|---|---|---|---|---|---|---|
| A | G | 12 | A | 0.3 | -0.563 | Destabilizing |
| A | L | 13 | A | 3.9 | -1.919 | Destabilizing |
| A | V | 14 | A | 7.7 | -1.653 | Destabilizing |
| A | N | 15 | A | 0.2 | -1.571 | Destabilizing |
| F | G | 12 | A | 0.3 | -0.464 | Destabilizing |
| F | L | 13 | A | 3.9 | -2.15 | Highly destabilizing |
| F | V | 14 | A | 7.7 | -2.071 | Highly destabilizing |
| F | N | 15 | A | 0.2 | -0.122 | Destabilizing |

### Supplementary Table 6 | HDX-MS data statistics

| | mKu | mKu + dsDNA | mKu + shDNA |
|---|---|---|---|
| ligand | - | linear dsDNA 40bp | Hairpin DNA (34 & 21 bp) |
| [mKu] (μM) | 15 | 15 | 15 |
| [DNA] (μM) | 15 | 15 | 15 |
| HDX buffer | 5mM K2HPO4, 5mM KH2PO4 pH 7 | | |
| HDX time | 0.1, 1, 10 & 100 min at 25°C | | |
| HDX control samples | mKu | | |
| Significance cut-offs | ΔHDX > 0.5 Da and *P*-value < 0.01in Welch's *t*-test (*n*=3) | | |
| No. of peptides | 208 | 208 | 208 |
| Sequence coverage (%) | 93.22 | 93.22 | 93.22 |
| Average peptide length | 11.02 | 11.02 | 11.02 |
| Peptide redundancy | 8.34 | 8.34 | 8.34 |
| Technical replicates | 3 | 3 | 3 |
| Mean SD | 0.087 | 0.089 | 0.090 |
| Data availability | PRIDE: PXD060776 | | |
